# Optimising age-specific insulin signalling to slow down reproductive ageing increases fitness in different environments

**DOI:** 10.1101/2023.12.22.573079

**Authors:** Zahida Sultanova, Aykut Shen, Katarzyna Hencel, Hanne Carlsson, Zoe Crighton, Daniel Clifton, Alper Akay, Alexei A. Maklakov

**Affiliations:** School of Biological Sciences, University of East Anglia, Norwich Research Park, Norwich NR4 7TU, UK

**Keywords:** Ageing, Life-History Evolution, Trade-Offs, Developmental Theory of Ageing

## Abstract

The developmental theory of ageing proposes that age-specific decline in the force of natural selection results in suboptimal levels of gene expression in adulthood, leading to functional senescence. This theory explicitly predicts that optimising gene expression in adulthood can ameliorate functional senescence and improve fitness. Reduced insulin/IGF-1 signalling (rIIS) extends the reproductive lifespan of *Caenorhabditis elegans* at the cost of reduced reproduction. Here, we show that adulthood-only rIIS improves late-life reproduction without any detrimental effects on other life-history traits in both benign and stressful conditions. Remarkably, we show that rIIS additively extends late-life reproduction and lifespan when animals are exposed to a fluctuating food environment – intermittent fasting (IF) – resulting in reduced food intake in early adulthood. Full factorial genome-wide RNA-Seq across the life course demonstrated that IF and rIIS modulate the age-specific expression of pro-longevity genes. IF, rIIS and combined IF + rIIS treatment downregulated genes involved in peptide metabolism in early life and differentially regulated immunity genes in later life. Importantly, combined IF + rIIS treatment uniquely regulated a large cluster of genes in mid-life that are associated with immune response. These results suggest that optimising gene expression in adulthood can decelerate reproductive ageing and increase fitness.

## Introduction

The evolutionary theory of ageing maintains that ageing evolves because the force of natural selection declines with age (*1*, *2*). The decline in selection gradients with age results in the accumulation of deleterious alleles whose effects on organismal performance are concentrated in late-life (mutation accumulation, MA) (*2*) and favours alleles whose effects on fitness are positive in early life despite being detrimental in late life (antagonistic pleiotropy, AP) (*3*). On a physiological level, there are two main proximate theories that aim to explain the evolution of ageing – the ‘disposable soma’ theory (DST) and the developmental theory of ageing (DTA)(*4*). The DST argues that ageing evolves via resource allocation trade-off between investment into somatic maintenance, growth, and reproduction (*5–7*). The DTA proposes that ageing evolves because selection fails to optimise age-specific gene expression, resulting in suboptimal physiology in late life (*4*, *8–11*). While the DST falls under the umbrella of AP, the DTA allows for the evolution of ageing via MA and AP alleles (*4*).

DST and DTA make unique testable predictions that can be used in empirical studies to advance our understanding of the evolutionary physiology of ageing (*4*). Specifically, the DTA predicts that optimising age-specific gene expression using experimental approaches can postpone or slow down age-related deterioration and improve fitness, while the DST predicts that diverting resources to improve late-life performance will come at the cost of early-life performance. Here we focused on a well-established system of insulin signalling-regulated life-history in *Caenorhabditis elegans* nematodes to ask whether downregulation of insulin/insulin-like signalling (IIS) in adulthood can slow down reproductive ageing without costs to other life-history traits, and, therefore, improve reproductive fitness.

Previous work established that reduced IIS extends reproductive lifespan in *C. elegans,* but such extension was accompanied by a marked reduction in overall reproduction (*12–15*) suggesting a genetic trade-off. It is important to emphasise that such genetic trade-off does not imply direct reallocation of resources between somatic maintenance and reproduction(*13*). Furthermore, while reduced IIS improves reproductive ageing in benign environments, it is not clear whether such effects will manifest in more natural conditions when the organisms are exposed to temporary food shortages. Interestingly, different forms of food shortages, such as continuous dietary restriction (reduced intake of nutrients without malnutrition) or intermittent fasting (IF) (aka time-restricted fasting) also can extend reproductive lifespan (*16*, *17*). However, it is not clear whether such effects are underpinned by the same molecular pathways. For example, it has been suggested that different forms of fasting increase lifespan via different molecular signalling pathways (*18*, *19*) and the same can apply to reproductive ageing.

IF is a form of dietary restriction (reduced nutrient intake without malnutrition) which involves cycling periods of fasting and food consumption (*20*). Different IF regimes have been shown to extend lifespan in model organisms and reduce age-related pathologies (*21*– *26*). IF is hypothesised to act via inhibiting insulin/IGF-1 signalling (IIS) in *C. elegans,* and previous work suggested that IF does not markedly increase lifespan in *daf-2* mutants (*23*, *26*). This could be in part due to the behaviour of the DAF-16 transcription factor, which translocates from the cytoplasm to the nucleus in response to fasting but relocates back to the cytoplasm under prolonged food shortage (over 24 hours) (*27*). Therefore, long-term fasting treatments likely bypass IIS and act via different pathways, including DAF-16 independent pathways such as AMPK (*18*, *19*, *28*). Similarly, it is possible that different IF regimes also act via different pathways and vary in their effect on late-life health (*18*).

Our goal was to establish whether adulthood-only downregulation of IIS can slow down reproductive ageing and improve organismal fitness under both ad libitum and limited resources. We found that reduced IIS not only improves reproductive ageing to increase lifetime reproductive success in different environments, but the benefit is particularly strong when resources are limited. Then, we investigated the impact of IF and rIIS on age-specific nuclear localisation of DAF-16 transcription factor and found that combined IF + rIIS treatment has a stronger effect across the life course because different treatments are more effective at different ages. IF leads to DAF-16 nuclear translocation during early-adulthood while rIIS causes a more moderate DAF-16 nuclear localisation during both early and late-adulthood. Therefore, when IF and rIIS are combined, this results in a more pronounced DAF-16 nuclear localisation from early to late-adulthood. Finally, we investigated the effects of both treatments on genome-wide gene expression in full-factorial design across three different ages (young, mid-life and old) and found that both similar and distinct physiological processes are modulated by early-adulthood IF and rIIS treatments at different ages. Early in life, shared mechanisms, such as suppressing translation, were activated in IF, rIIS and their combination. In contrast, late in life, more distinct mechanisms, such as the ones involved in immunity, were differentially regulated by IF, rIIS and their combination.

## Results

### Early-adulthood IF and rIIS additively increase survival and slow down reproductive ageing

To understand whether rIIS and IF work through similar mechanisms as previously suggested, we explored the effects of rIIS in combination with IF. Considering that the timing of dietary interventions matters, we examined how early and late-adulthood IF influenced survival (Figs 1, 2 and S1). We found that rIIS and early-adulthood IF additively increased survival in the wild-type N2 strain (Fig. 2A).

**Fig. 1.**
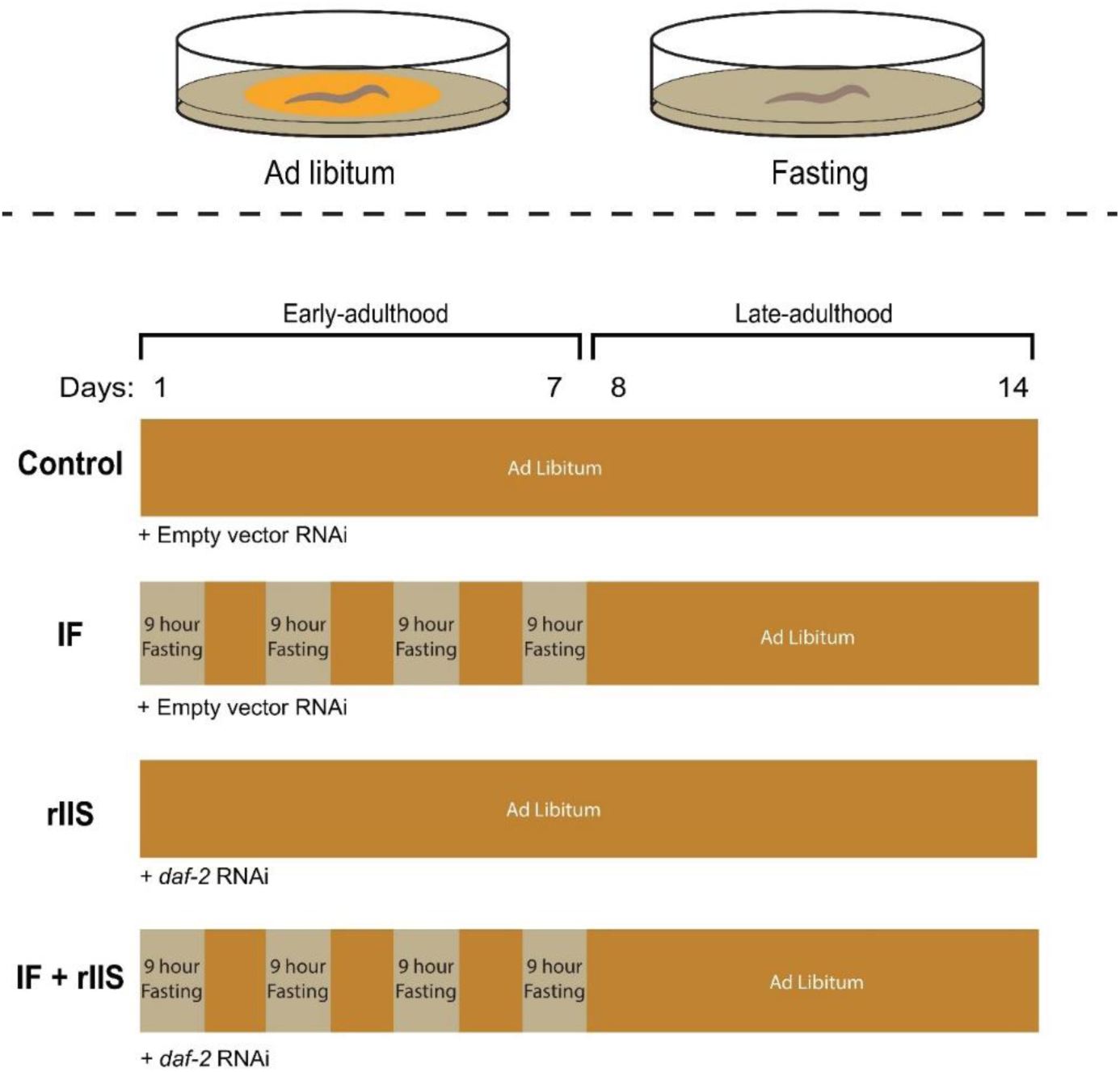
Schematic illustration of the experimental treatments. Late L4 stage worms were subjected to four different experimental treatments (Control, IF, rIIS and combined). Early-adulthood intermittent fasting involved fasting for 9 hours on alternate days during the initial week of lifespan. Late-adulthood intermittent fasting involved fasting for 9 hours on alternate days during the second week of lifespan. Control and IF treatments were fed with *E. coli* that produces empty vector RNAi; rIIS and combined treatments were fed with *E. coli* that produces *daf-2* RNAi.

**Fig. 2.**
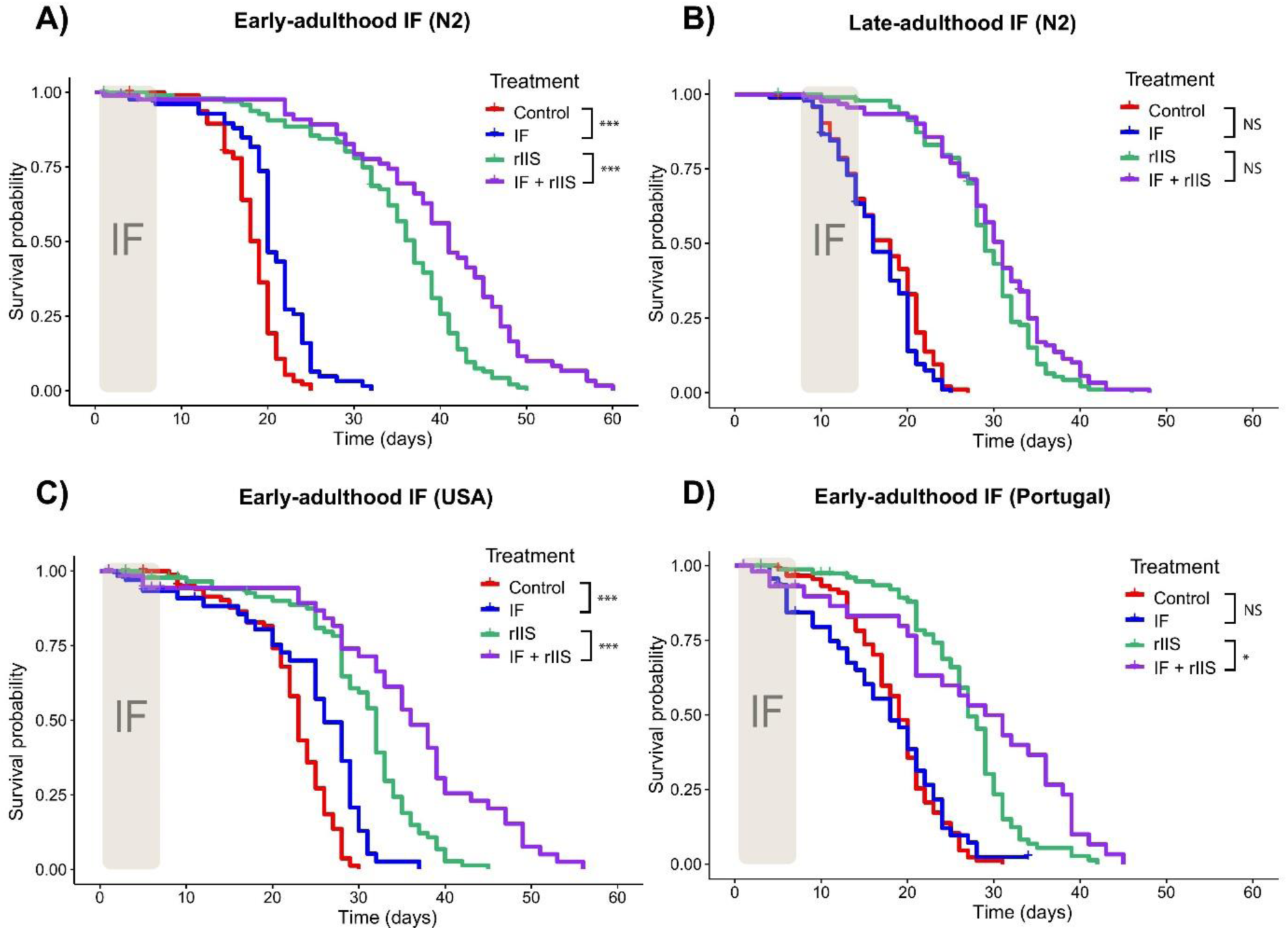
Effect of IF and rIIS on survival. Effect of early and late-adulthood IF on survival of N2 worms with and without rIIS (A and B, respectively). Effect of early-adulthood IF on survival of USA and Portugal strains with and without rIIS (C and D, respectively).Significance in the divergence of survival curves was assessed using log-rank tests, with statistical significance represented as follows: NS indicates P > 0.05, * indicates P ≤ 0.05, ** indicates P ≤ 0.01, and *** indicates P ≤ 0.001.

In contrast, late-adulthood IF did not affect survival, regardless of whether rIIS was present or not (Fig. 2B). We also assessed the impact of rIIS and early-adulthood IF on various strains of *C. elegans* to evaluate the generalisability of our findings beyond N2, the commonly used laboratory strain. For USA strain, early-adulthood IF increased survival both in the absence and presence of rIIS (Fig. 2C). For Portugal strain, there was no effect of early-adulthood IF on the survival of control worms but it further increased survival in the presence of rIIS (Fig. 2D). In summary, our findings showed that rIIS increases survival when combined with early-adulthood IF and this was consistent across different worm strains.

Knowing that various dietary restriction types and rIIS mutants can reduce reproduction, we investigated how IF and rIIS affect self-fertilised reproduction. IF significantly reduced reproduction (Fig. 3A, IF: χ² = 76.41, df = 1, p < 0.001), whereas rIIS had no effect on reproduction in the N2 strain (rIIS: χ² = 0.65, df = 1, p = 0.421; IF x rIIS interaction: χ² = 2.41, df = 1, p = 0.121). These effects did not change when matricides were censored and were similar in two other *C. elegans* strains (Fig. S2). In brief, we showed that, in contrast to intermittent fasting, reduced IIS does not decrease reproduction during its application.

**Fig. 3.**
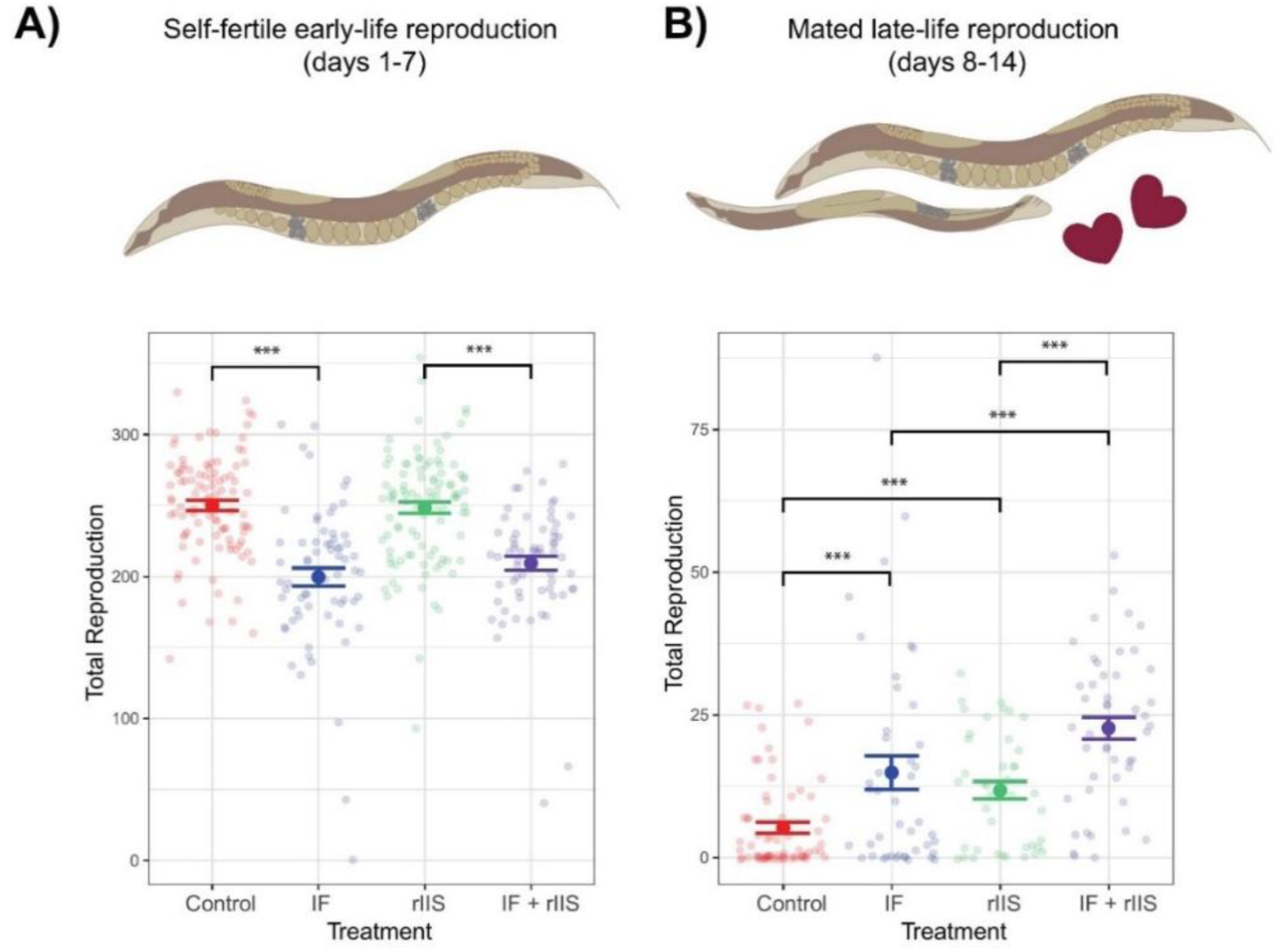
Effect of IF and rIIS on reproduction. For early-life self-fertile reproduction (A), differences between treatments were assessed by Tukey’s HSD test with Benforini-Hochberg correction for multiple testing. Early-life self-reproduction data presented here is also discussed in Duxbury *et al*., (2022), highlighting its relevance in a different context (*29*). For late-life mated reproduction (B), differences between treatments were assessed by Dunn test with Benforini-Hochberg correction for multiple testing. NS indicates P > 0.05, * indicates P ≤ 0.05, ** indicates P ≤ 0.01, and *** indicates P ≤ 0.001.

To test whether IF and rIIS slow down reproductive ageing, we explored the effects of IF and rIIS on late-life mated reproduction. Our study revealed that both IF and rIIS independently led to increased late-life reproduction (Fig. 3B, IF: χ² = 16.5049, df = 1, p < 0.001; rIIS: χ² = 6.9117, df = 1, p = 0.009; IF x rIIS interaction: χ² = 0.0122, df = 1, p = 0.912). When IF and rIIS were combined, late-life reproduction was higher than all other treatments (Fig. 3B and S3). Altogether, these data emphasize that IF and rIIS additively delay reproductive ageing.

Finally, we tested whether IF and rIIS have effects that pass down to the next generation by looking at the survival and reproduction of offspring produced later in life. Our findings show that neither IF nor rIIS changed how well these offspring survived (Fig. S4). Similarly, there was no effect of IF or rIIS on offspring lifetime reproductive success (Fig. S5, IF: χ² = 0.8584, df = 1, p = 0.354; rIIS: χ² = 2.5170, df = 1, p = 0.113; IF x rIIS interaction: χ² = 0.6178, df = 1, p = 0.432).

### Both IF and rIIS enhance DAF-16 nuclear localisation but in an age-specific manner

To understand the potential mechanisms underlying the effects of IF and rIIS on ageing, we explored the nuclear localisation of the DAF-16 transcription factor. DAF-16 activates longevity genes when it moves into the nucleus (*28*). We observed that IF significantly increased DAF-16 nuclear localisation during early-adulthood (Fig. 4A, B), but this effect diminished gradually with advancing age (Fig. 4C, D). On the other hand, rIIS also caused DAF-16 to move into the nucleus, but this effect was weaker compared to IF during early-adulthood (Fig. 4A, B). Interestingly, unlike IF, rIIS continued to enhance DAF-16’s localisation into the nucleus as worms grew older (Fig. 4C, D). These varying effects at different ages led to a significant interaction between IF and rIIS (χ² = 5.12, df = 1, p = 0.02) with marginally nonsignificant interactions between rIIS and age (χ² = 3.18, df = 1, p = 0.07) and between IF and age (χ² = 3.10, df = 1, p = 0.08). Thanks to the early-life effect of IF and the sustained impact of rIIS during both early and late life, the combined treatment benefited from higher DAF-16 nuclear localisation throughout the lifespan (Fig. 4).

**Fig. 4.**
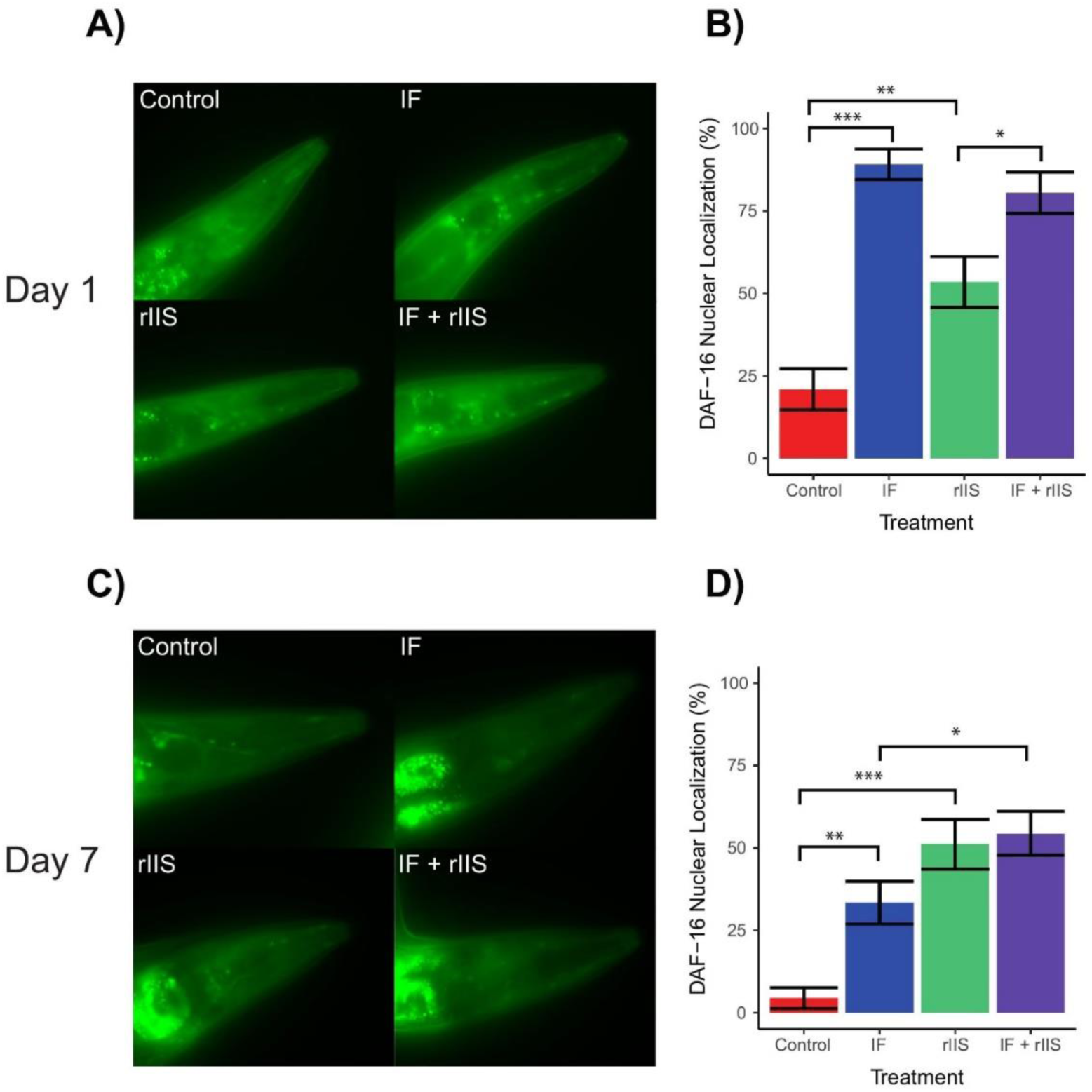
Effect of IF and rIIS on DAF-16 nuclear localisation. Photographs and corresponding bar plots illustrating the effect of experimental treatments on the nuclear localisation of the DAF-16 transcription factor tagged with GFP, assessed on day 1 (panels A and B) and day 7 (panels C and D). The differences between treatments were evaluated using chi-square tests with Benforini-Hochberg correction for multiple testing. NS indicates P > 0.05, * indicates P ≤ 0.05, ** indicates P ≤ 0.01, and *** indicates P ≤ 0.001.

### IF, rIIS and their combined treatment exert age-dependent effects on gene expression

We analysed gene expression changes in worms subjected to IF, rIIS or both on day 1, day 7 and day 13 by sequencing polyA(+) RNA in triplicate for each condition and time point. The principal component analysis (PCA) of samples shows that the largest variation in gene expression changes is due to the ageing of animals, as PC1 can separate all Day 1 animals from Day 7 and Day 13 (Fig. S6). Next, we calculated the differential gene expression changes in worms exposed to IF, rIIS and combined treatments compared to control worms (Fig. 5A). The largest gene expression changes are observed in day 1 animals, and the number of differentially expressed genes between conditions go down as the animals age for day 7 and day 13 samples. Moreover, while rIIS treatment resulted in the highest number of differentially expressed genes in the early life, the combined treatment led to a greater number of differentially expressed genes at more advanced ages (Fig. 5A).

**Fig. 5.**
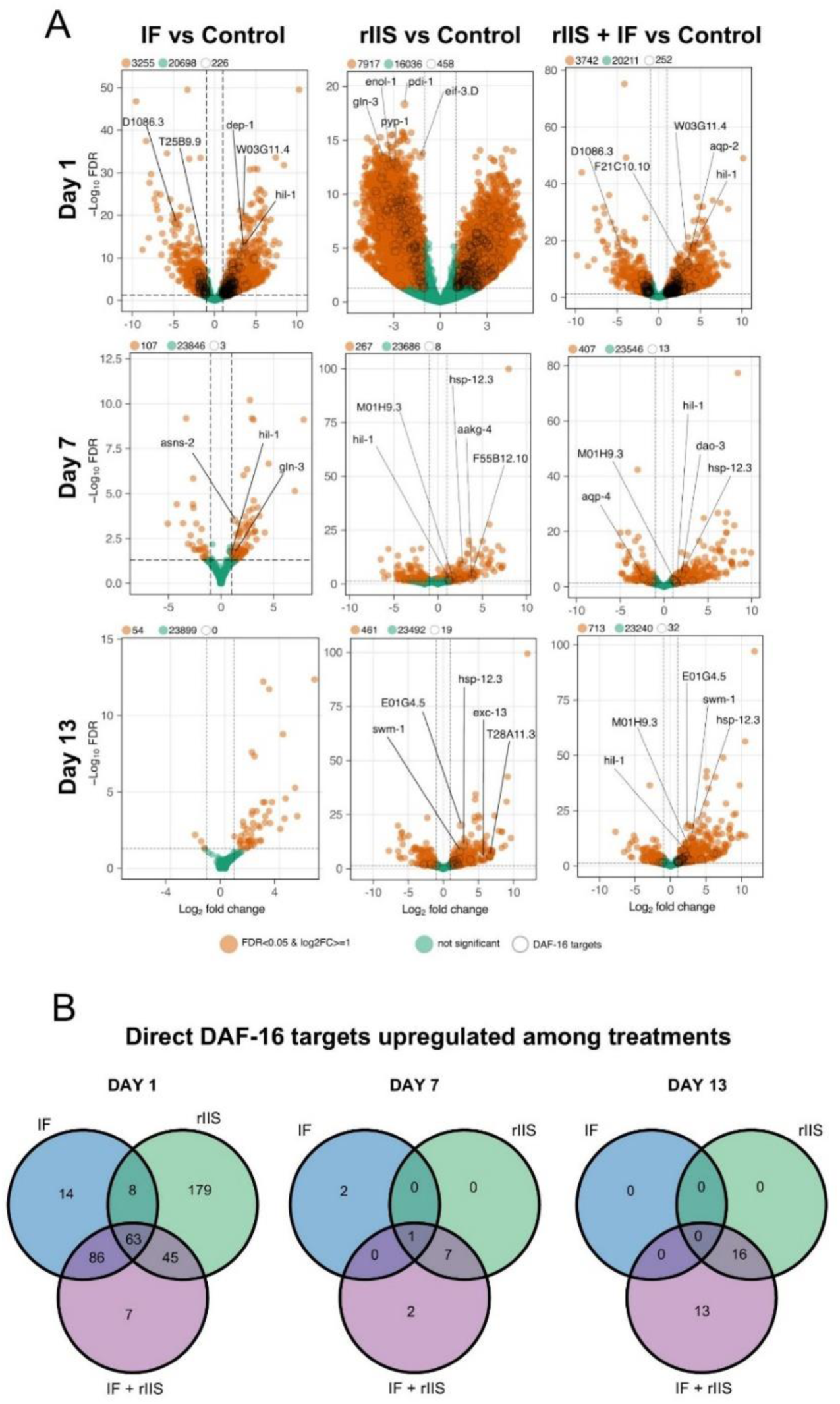
Effect of IF and rIIS on age-specific gene expression. Volcano plots comparing differentially expressed genes in combined treatment compared to control. The top 5 direct DAF-16 targets that were differentially expressed in each treatment with respect to statistical significance were shown on each plot. DAF-16 direct targets were taken from a previous study that identified them by using chromatin profiling by DNA adenine methyltransferase identification, DamID (*30*) (A). Venn diagrams showing the number of direct DAF-16 targets that are upregulated across IF, rIIS and combined treatments compared to control (*30*) (B).

Next, to understand how DAF-16 activation contributes to the differential gene expression across treatments and time points, we analysed the presence of previously identified direct targets of DAF-16 (*30*). Direct DAF-16 targets behaved similar to the total number of differentially expressed genes. During early adulthood (day 1), most of the direct DAF-16 target genes were activated by rIIS. As the worms aged (days 7 and 13), rIIS alone was not enough and the combination of rIIS and IF was necessary to activate many DAF-16 direct targets (Fig. 5B).

### IF, rIIS, and their combination have age-specific effects on diverse biological functions later in life, particularly those related to biosynthesis and immune response

We conducted a Gene Ontology analysis to explore the functions of genes differentially expressed with age across different treatments. First, we looked at genes differentially expressed in the combined rIIS + IF treatment compared to the control to understand which genes are affected by the combination of rIIS with IF (Fig. 6, see Figs. S7 & S8 for IF and rIIS treatments). During early adulthood (day 1), the combined treatment enhanced the expression of genes involved in developmental processes and simultaneously reduced the expression of genes associated with peptide metabolism (Fig. 6A). Conversely, in late adulthood (days 7 and 13), distinct groups of genes associated with immune response exhibited varied expression patterns, some upregulated, and others downregulated (Fig. 6B and C).

**Fig. 6.**
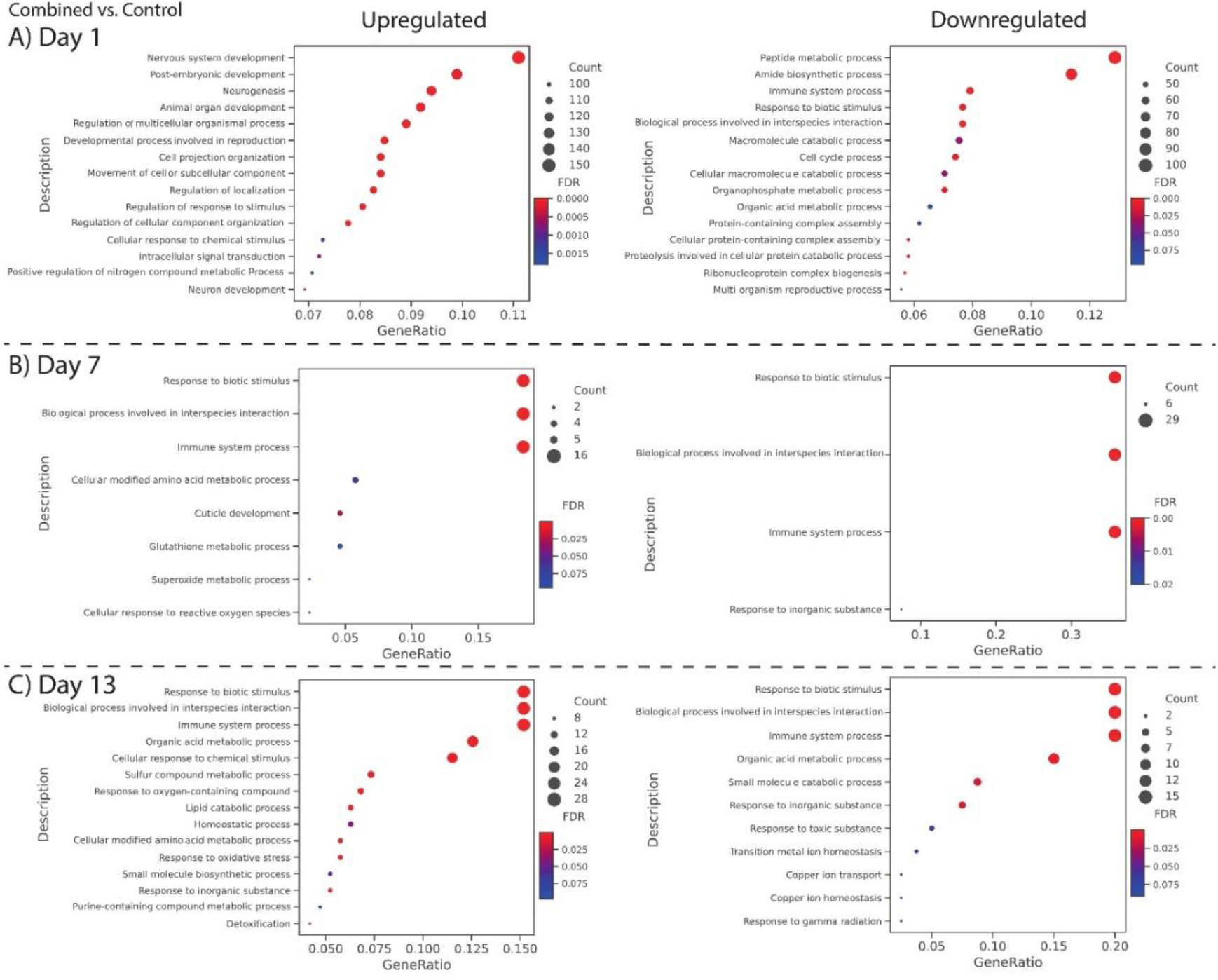
Gene Ontology analysis of differentially expressed genes in the combined treatment. GO term analyses showing the functions of genes up and down-regulated by the combined treatment compared to control across day 1 (A), day 7 (B) and day 13 (C).

To understand the shared and unique mechanisms between IF and rIIS, we used Venn diagrams for Gene Ontology (Figs. 7-9). On day 1, all treatments upregulated genes involved in developmental processes. Moreover, rIIS uniquely upregulated 55 % of differentially expressed genes. These genes were related to processes such as DNA and RNA metabolism (Fig. 7A). On the other hand, all treatments downregulated genes involved in processes like peptide metabolism, ATP metabolic process and Ribonucleoprotein complex biogenesis. In addition to that, IF uniquely downregulated genes involved in carbohydrate biosynthesis. 56 % of differentially regulated genes were uniquely downregulated by rIIS, involved in processes such as neuropeptide signalling, tricarboxylic acid cycle and actin filament regulation (Fig. 7B). In summary, during early adulthood, all treatments upregulated genes involved in developmental processes while downregulating genes associated with biosynthesis.

**Fig. 7.**
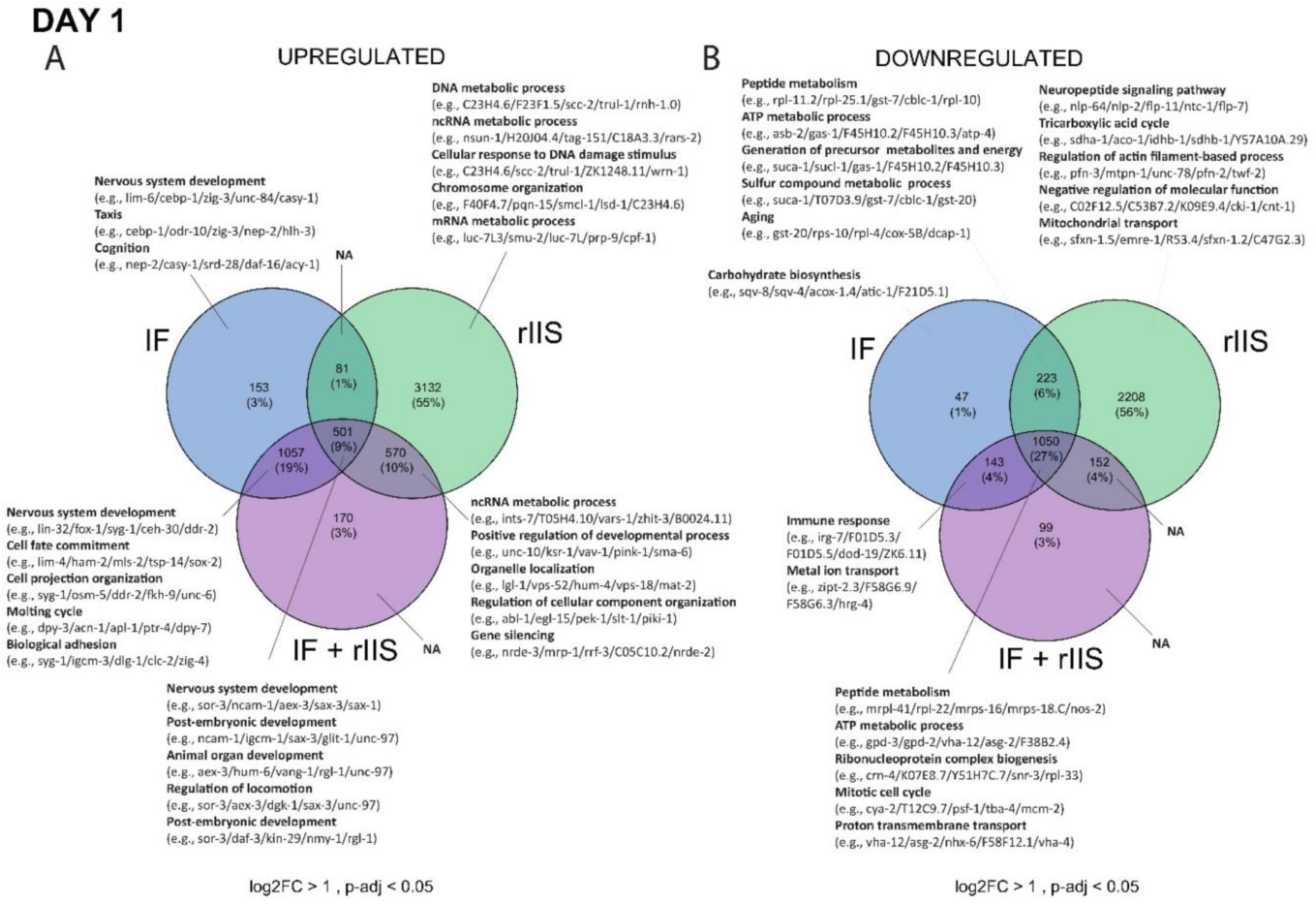
Gene Ontology analysis of differentially expressed genes across IF, rIIS, and combined treatments on day 1. GO term analyses showing the functions of genes up-regulated (A) and down-regulated (B) by IF, rIIS, and combined treatments compared to the control on day 1.

On day 7, IF, rIIS and their combination differentially regulated several immune genes (Fig. 8). Although many immune-related genes were shared between rIIS and the combined treatment, there were others that were only expressed when rIIS was combined with IF. Specifically, the combined and IF treatments together upregulated immune genes *cnc-1*, *cnc-5* and *clec-52*, which are known to function through pathways beyond IIS, such as TGF-β and p38 MAPK pathways (*31*, *32*) (Fig. 8A). Likewise, the combined treatment also uniquely downregulated a substantial number of genes related to immunity (Fig. 8B). Intriguingly, among them, *cdr-4*, previously reported to be upregulated by rIIS (*33*), exhibited the opposite response in the combined treatment. In brief, the combination of rIIS and IF shares some immune genes with rIIS, but it also influences other genes not connected to insulin signalling, potentially through the effects of intermittent fasting.

**Fig. 8.**
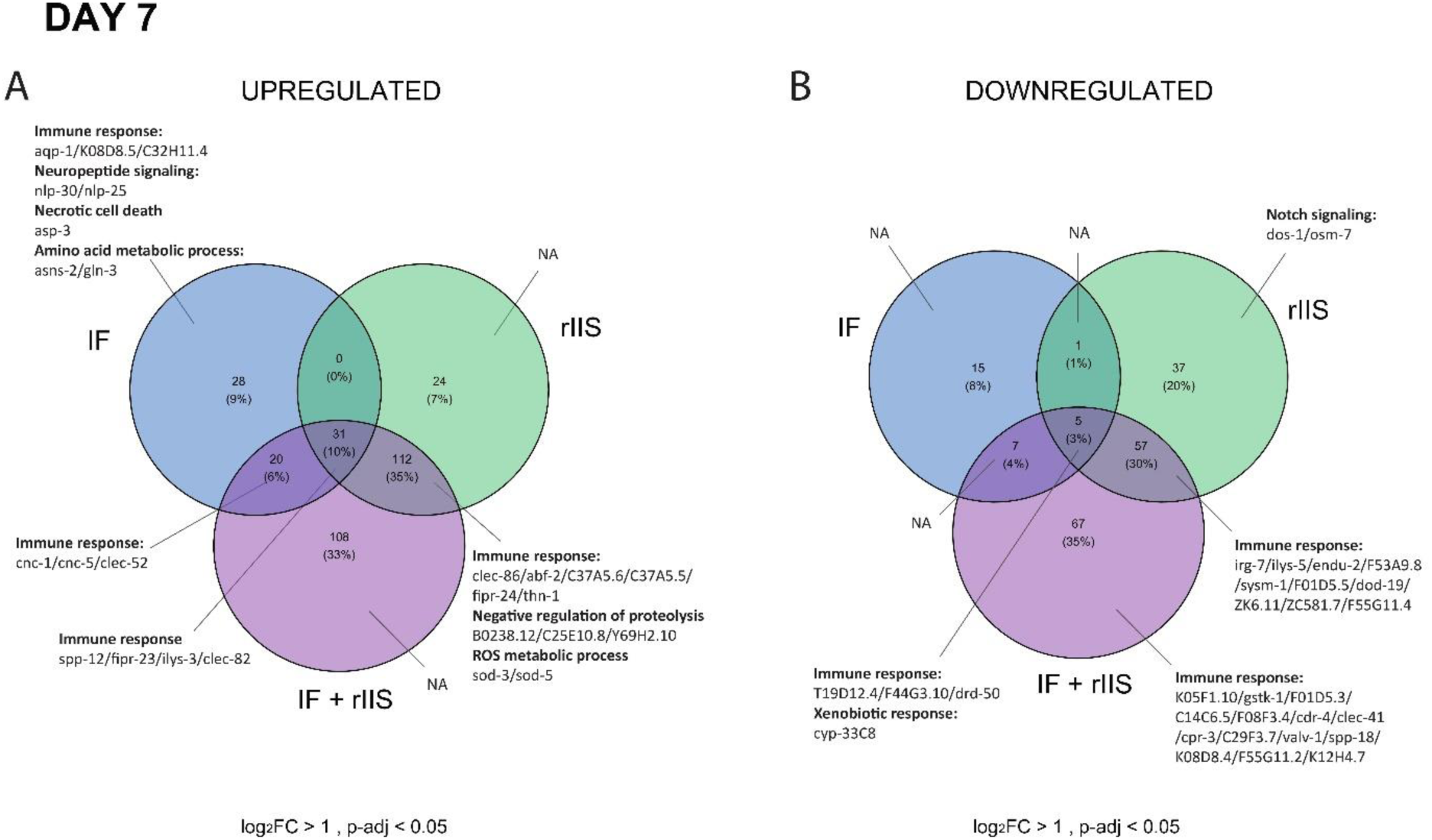
Gene Ontology analysis of differentially expressed genes across IF, rIIS and combined treatments on day 7. GO term analyses showing the functions of genes up-regulated (A) and down-regulated (B) by IF, rIIS and combined treatments compared to the control on day 7.

**Fig. 9.**
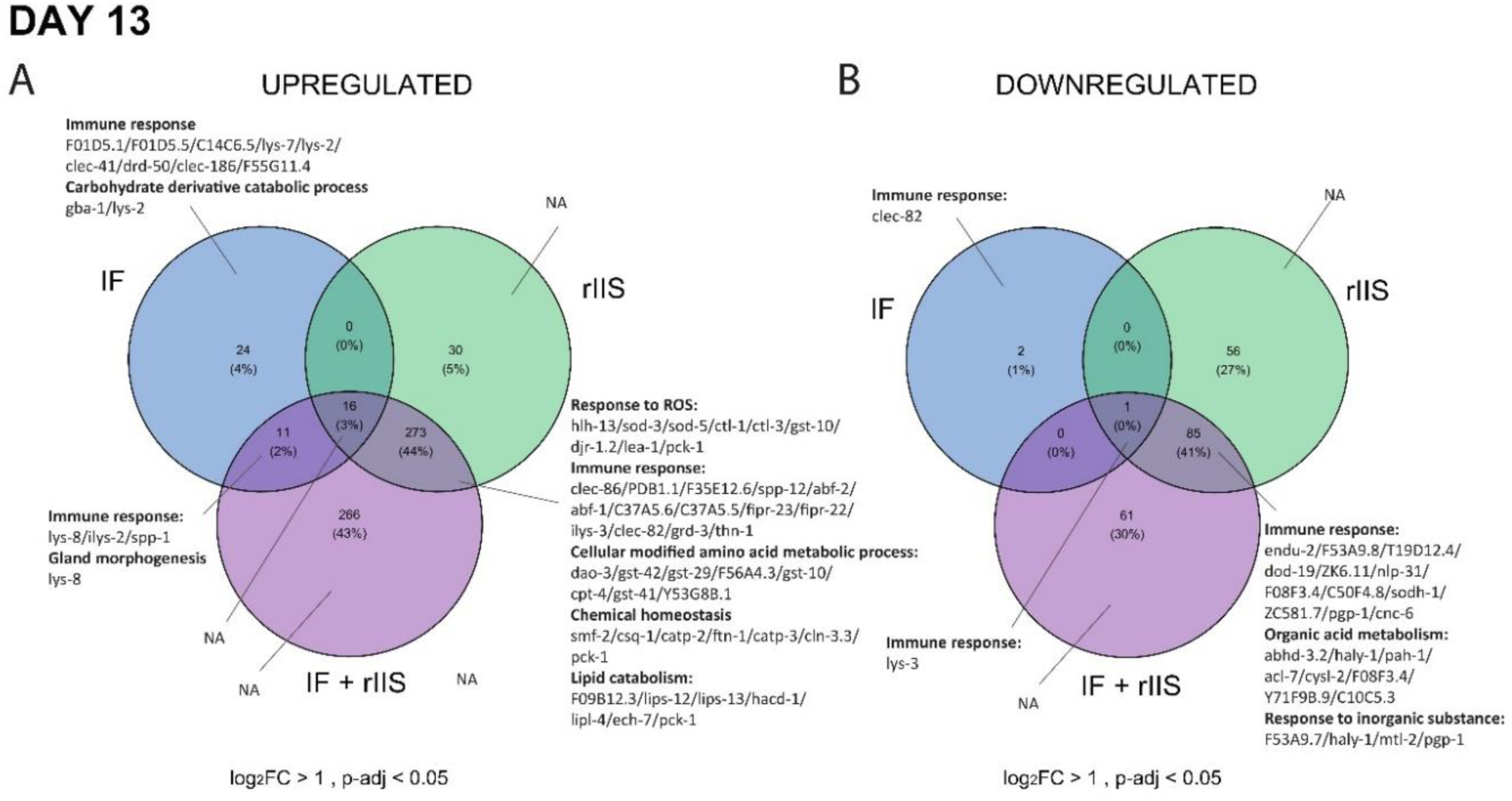
Gene Ontology analysis of differentially expressed genes across IF, rIIS and combined treatments on day 13. GO term analyses showing the functions of genes up-regulated (A) and down-regulated (B) by IF, rIIS and combined treatments compared to control on day 13.

On day 13, similar to the patterns observed on day 7, the combination of rIIS and IF exhibited differential regulation of various immune genes, often together with rIIS (Fig. 9). rIIS and the combined treatment both upregulated genes involved in several processes, including immune response, response to ROS and amino acid metabolism. On the other hand, the combined and IF treatments jointly upregulated immune genes *lys-8*, *spp-1* and *ilys-2* (Fig. 9A). Among them, *lys-8* has been reported to be influenced by IIS, as well as TGF-β and p38 MAPK pathways (*34*). Interestingly, both *spp-1* and *ilys-2* were found to be downregulated by rIIS, which is the opposite direction of how combined treatment and IF regulates them (*33*). So, only *lys-8* was regulated in the same direction as rIIS. rIIS and the combined treatment both downregulated mechanisms associated with immune response, organic acid metabolism and response to inorganic substances (Fig. 9B). Intriguingly, no genes were downregulated by IF and the combined treatment together. In short, at older ages, the combination of rIIS and IF regulates a substantial number of processes similar to rIIS alone, while some processes are also influenced by IF.

## Discussion

We found that adulthood-only reduced insulin signalling (rIIS) via *daf-2* knockdown slows down reproductive ageing and, in doing so, increases lifetime reproductive success in *C. elegans* hermaphrodites because early life reproduction via selfing was unaffected. This effect was particularly pronounced when animals experienced food shortage in the form of intermittent fasting (IF). By examining the nuclear localisation of DAF-16 across different ages, we showed that the combined effects of early-adulthood IF and rIIS led to higher total nuclear localisation of this transcription factor, potentially maximizing its activity.

Specifically, DAF-16 localisation in the combined treatment was mainly driven by IF instead of rIIS in early life, while the opposite was true later in life. Genome-wide RNA sequencing showed that IF and rIIS shape gene expression in an age-dependent manner. The GO terms analyses suggest that genes involved in biosynthesis and immunity are potential candidates for explaining the additive effects of IF and rIIS on ageing. The reduction in biosynthesis during early adulthood at least partially aligns with the disposable soma theory (DST), suggesting a trade-off between growth and maintenance (resource allocation). In contrast, regulation of immunity genes later in life is broadly in line with the developmental theory of ageing (DTA), which posits that ageing results from suboptimal gene expression in late life due to the weakening of natural selection on biological function in adulthood.

Previous work shows a complex picture with respect to the suggested role of insulin signalling in reproduction in *C. elegans*. Different studies reported that while some *daf-2* alleles can result in sterility (*35*), others lead to the same (*14*, *36*) or extended reproductive lifespan (*35*, *37*) of selfing hermaphrodites compared to wild-type worms but with severely reduced reproductive rate and total brood sizes, especially in stressful and fluctuating temperatures (*35*). However, self-fertile *C. elegans* hermaphrodites are sperm limited (they produce around 300 sperm prior to maturation), and mating can substantially increase late reproduction and lead to higher lifetime reproductive success. Therefore, while some reproductive ageing in *C. elegans* hermaphrodites occurs prior to cessation of sperm, it is key to study reproductive ageing in mated hermaphrodites that have unlimited access to sperm while still producing oocytes. Mated *daf-2 (e1370)* hermaphrodites indeed can have increased reproductive lifespan but reduced rate of reproduction and low lifetime reproductive success compared to wild-type animals (*15*). Later work demonstrated that *daf-2* mutants have improved maintenance of oocyte quality with age (*12*, *38*). Because early-life reproduction is more important for fitness than late-life reproduction, given *C. elegans’* life history (*39*), this suggests a genetic trade-off or constraint that links reproductive lifespan extension to reduced fitness. It is important to emphasise that such a trade-off is unlikely to be resource-dependent because neither reduction nor increase in early-life offspring production has a direct effect on reproductive ageing (*12*, *13*, *15*). Finally, later work suggested that mated *daf-2 (e1370)* hermaphrodites may actually suffer from reduced reproductive lifespan and reduced overall reproduction (*40*). The discrepancy between different studies likely results from the timing of mating during life course and differences in animal husbandry, but it is clear that lifelong reduction in IIS in *daf-2* mutants is likely to carry substantial fitness costs, even if it has positive effects on reproductive ageing under certain conditions.

While we found that adulthood-only *daf-2* RNAi improves late post-mating reproduction without negative effects on the early self-fertile stage together with increased longevity, it still remains possible that there is a trade-off between slower reproductive ageing and offspring quality. We have previously shown that adulthood-only *daf-2* RNAi improves offspring quality of self-fertile hermaphrodites in a benign environment (*41*), has no effect on offspring quality in fluctuating stressful environments (*42*) and has overall positive effects on fitness and lineage survival in a long-term multigenerational scenario, which mimicked the evolution of age-specific modifier gene that reduces adulthood-only expression of daf-2 in every generation (*29*). Here, we found that offspring produced late in life by *daf-2* RNAi mated hermaphrodites had similar lifetime reproductive success and survival compared to wild-type worms. Overall, these results suggest that *daf-2* RNAi-mediated improvement in reproductive ageing is both general across benign and stressful environments and does not trade-off with offspring quality.

Interestingly, we found that IF during early-adulthood, but not late-adulthood, increases survival. This result aligns with previous work in *D. melanogaster* fruit flies, suggesting that IF has a greater impact on survival when applied early, but not late, in life (*22*, *25*). In fruit flies, 5 days fasting-2 days eating IF regimen increased lifespan when applied during early or mid-adulthood. However, when applied during late-adulthood, it shortened lifespan (*22*). Similarly, 18 hours fasting-6 hours eating IF regimen also increased lifespan in fruit flies when applied during early, but not late, adulthood (*25*). Our findings in worms further support the importance of the timing of the treatment, with early-adulthood being more beneficial.

Crucially, early-adulthood IF increased survival both in the presence and absence of rIIS, suggesting that IF may be working at least in part through a different route than IIS. Although early or late-adulthood IF was not previously compared in *C. elegans*, it was shown that applying IF throughout the entire lifespan increases longevity (*23*, *26*). In these studies, a 2-day fasting-2-day eating regimen increased survival in *C. elegans* through an IIS-dependent mechanism, contrasting with our findings (*23*, *26*). Our IF regimen differs from previous studies, which used a 2-day intermittent fasting and eating cycle (*26*), whereas in our experiment, we applied 9 hours of fasting on alternate days. In fact, our IF regimen is perhaps more similar to the time-restricted feeding regimen used in fruit flies, which was suggested to work independently of inhibition of insulin-like signalling (*25*).

To gain further insights into the mechanisms underlying our results, we first explored DAF-16 nuclear localisation. DAF-16 is a key transcription factor which acts downstream of nutrient-sensing signalling and germline signalling pathways (*28*). As a response to signals related to food availability or germline maintenance, it enters the nucleus and transcriptionally activates pro-longevity genes (*28*). In our experiment, DAF-16 is expected to remain in the cytoplasm in the presence of food and when the IIS pathway is active (control). However, when food is scarce (IF treatment), when the IIS is inactivated (rIIS treatment), or both (combined treatment), DAF-16 is expected to relocate to the nucleus (*19*). Our findings indicate that in early-adulthood, fasting causes DAF-16 to enter the nucleus more than rIIS. Conversely, fasting is no longer that effective in later adulthood, while rIIS worms maintain the same DAF-16 nuclear localisation as observed in their youth. DAF-16 has been shown to rapidly respond to changing food availability by moving into the nucleus shortly after fasting initiation and returning to the cytoplasm upon resuming a normal diet (*19*). Hence, in our IF treatment, DAF-16 is expected to be in the nucleus only during fasting periods (9 hours on alternate days) and back in the cytoplasm during non-fasting periods. In the rIIS treatment, DAF-16 is expected to be in the nucleus constantly, at least during the first week of lifespan, because IIS is constantly reduced by RNAi treatment. In fact, deactivating the IIS pathway has been shown to cause DAF-16 to stay in the nucleus even until later ages, as late as day 11 (*27*). In the combined treatment, worms experience both intermittent nuclear localisation due to IF and the constant nuclear localisation due to rIIS, and benefit from both treatments. These findings support the hypothesis that IF and rIIS are modulating DAF-16 localisation in an age-dependent manner, leading to potentially increased DAF-16 activity in combined treatment. This is further supported by the fact that some direct DAF-16 targets are only expressed in the combined treatment at later ages.

To better understand the mechanisms driving the effects of the combined treatment, we explored the function of genes that were differentially expressed by IF, rIIS and their combination at different ages. Our findings revealed that IF and rIIS activate both similar and distinct mechanisms depending on the age of the worms. For example, during early adulthood, all treatments downregulate numerous genes involved in biosynthesis, a change known to promote longevity (*43*, *44*). Likewise, in later adulthood, they all downregulate a gene involved in the xenobiotic response*, cyp-33C8*, whose downregulation increases survival under oxygen deprivation (*45*). This implies that similar mechanisms could contribute to extending reproductive ageing and lifespan across IF, rIIS, and combined treatments, particularly early in life. Still, the level of downregulation is important because decreasing the synthesis of proteins and carbohydrates can be harmful to reproduction (*46*, *47*). In fact, we observed that IF uniquely downregulates genes involved in carbohydrate biosynthesis early in life. Differential regulation of biosynthetic processes can be one of the reasons why rIIS had no effect on early reproduction while IF was detrimental.

However, later in life, the combined treatment uniquely combines distinct and shared immune mechanisms originating from both IF and rIIS. Notably, numerous immune genes upregulated in the combined treatment together with IF are not related to the IIS pathway, indicating that IF can trigger additional immune responses through alternative mechanisms (*31*, *32*). Although the immune response is crucial for survival and reproduction, excessive production of antimicrobial peptides can actually harm the host due to elevated inflammation (*48*). For example, increased expression of some of the immune genes can lead to poor reproductive success (*49*). Therefore, finetuning the age-specific expression of immunity genes has an important role in regulating lifespan and reproduction. In fruit flies, selection for late-life reproduction and longevity led to the enrichment of immunity genes, suggesting that age-specific regulation of these genes contributes to extended lifespan and reproductive-span (*50*). The simultaneous control of various immune response pathways by the combined treatment could provide insight into its additive effect on survival and late-life reproduction.

Nevertheless, it is important to note that the worms in our study were kept in clean environments without any infections. In fact, the downregulation of certain genes involved in immunity by the combined treatment could be detrimental and pose challenges in the presence of infectious threats. For example, we found that *lys-3*, a lysozyme induced in response to pathogenic bacteria (*51*), is consistently downregulated in all three treatments during late adulthood, despite evidence that its overexpression can extend lifespan in *C. elegans* (*52*). Therefore, it becomes evident that not all mechanisms regulated by the combined treatment are universally beneficial. Finetuning immune response genes is indeed challenging but it could be a powerful way to slow down ageing effectively.

Our study suggests that reduced insulin signalling in adulthood ameliorates reproductive ageing in benign and food-limited environments without costs to reproductive rate or offspring quality. Notably, the benefits of adulthood-only rIIS to late-life reproduction and survival become more apparent when animals experience fluctuating food shortages in the form of IF. Specifically, we observed that early-adulthood IF and adulthood-only rIIS additively enhance both survival and late-life reproduction. The dynamic regulation of DAF-16 nuclear localisation in an age-specific manner suggests a finely tuned mechanism through which IF and rIIS exert their effects. Our exploration of age-specific gene expression patterns suggests that both shared and unique regulatory pathways may contribute to the additive effects of IF and rIIS on life-history traits. These results are in line with the hypothesis that reduction in the force of natural selection after reproductive maturity results in the evolution of suboptimal gene expression in adulthood, causing ageing (*4*, *9*).

It has been previously suggested that both DTA and DST-like processes can play a role in the evolution of ageing, and that these processes are not mutually exclusive (*4, 11*). The additive increase in lifespan and late-life reproduction when rIIS was combined with IF aligns with this suggestion because IF directly limits reproduction, and combining rIIS with IF to further slow down reproductive ageing comes at the cost of early-life reproduction. Further research is necessary to establish the relative contributions of resource allocation trade-offs and suboptimal age-specific gene expression to the evolution and expression of ageing across a broad range of taxa.

## Materials and Methods

### Experimental design

Three *C. elegans* wild-type strains: N2 (UK, Bristol), JT11398 (Portugal), EG4725 (USA) and the mutant strain: zIs356 [daf-16p::daf-16a/b::GFP + rol-6(su1006)] were obtained from Caenorhabditis Genetics Center (University of Minnesota, USA, funded by NIH Office of Research Infrastructure Programs, P40 OD010440) and stored at −80°C until use. Prior to the setup, the defrosted *C. elegans* strains were propagated for two generations on NGM agar supplemented with nystatin (100 μg/mL), streptomycin (100 μg/mL), and ampicillin (100 μg/mL) to prevent infection, following standard recipes (*53*). The agar plates were seeded with the antibiotic-resistant *Escherichia coli* bacterial strain OP50-1 (pUC4K, from J. Ewbank at the Centre d’Immunologie de Marseille-Luminy, France). To standardize parental age and eliminate any effects from defrosting, we bleached the eggs from the grandparents of experimental individuals before conducting the experiments.

To decrease the expression of the insulin/IGF-1 signalling receptor homolog, *daf-2*, during adulthood, we fed late-L4 larvae with *E. coli* bacteria that expressed *daf-2* double-stranded RNA. As a control, we used RNase-III deficient, IPTG-inducible HT115 *E. coli* bacteria containing an empty vector plasmid (L4440) (*41*, *54*). The RNAi clones were obtained from the Vidal feeding library (Source BioScience), which was created by M. Vidal lab at Harvard Medical School, USA. Prior to delivery, all clones were verified through sequencing.

To investigate the effects of intermittent fasting (IF) and reduced insulin signalling (rIIS) on *C. elegans* survival, early and late-life reproduction, we designed four distinct treatments (Fig. 1). **“Control”** treatment group consisted of worms kept on standard NGM agar plates with an *E. coli* lawn that produced empty vector RNAi. **“IF”** treatment group consisted of worms exposed to a novel IF regimen during one week of their lifespan by being fed with empty vector RNAi. In this IF regime, we kept worms on foodless plates for 9 hours on alternate days. These foodless plates were not seeded with *E. coli* and contained no peptone, a crucial ingredient for bacterial growth. **“rIIS”** treatment group included worms treated like control treatment but maintained on *daf-2* RNAi producing *E. coli* lawn. Finally, **“IF + rIIS (combined)”** treatment group included worms treated like IF treatment but kept on *daf-2* RNAi producing *E. coli* lawn. Thus, we had a fully factorial design with two different feeding treatments and two different RNAi treatments. This approach allowed us to examine the combined effects of IF and rIIS on survival and reproduction.

### Survival assays

To investigate the effects of IF and rIIS on worm survival, we explored how IF early and late in adulthood influences lifespan of worms in the absence and presence of rIIS. For early-adulthood IF, we kept the worms individually during their self-reproductive period. After reproduction ceased, we grouped the worms into cohorts of approximately 10 individuals. For late-adulthood IF, we always kept the worms in groups until they died. We monitored survival until the death of each worm, censoring any worms that were lost during the experiment.

### Reproductive assays

We monitored self-reproduction during the first week of each worm’s lifespan. For the Control and rIIS treatment groups, worms were transferred to new plates every 24 hours. For the IF and IF + rIIS treatment groups, worms were transferred to new plates twice on fasting days and every 24 hours on ad libitum (adlib) days. To ensure that the eggs laid during the 7-day reproductive period developed into larvae, we allowed the eggs to hatch and develop for two days. After this period, the hatched larvae were killed at 42°C for 3.5 hours and stored for counting later.

We monitored late-life reproduction by mating hermaphrodites with males after the self-reproductive period ended. Males were produced by exposing around 50 L4 hermaphrodites to 30°C heat shock for 5 hours on NGM plates and then incubating them at 20°C. On days 8 and 12, two males were paired with a single hermaphrodite for 24 hours, after which the males were discarded. The hermaphrodites were then transferred to fresh plates daily until the cessation of reproduction on day 14. Hermaphrodites were maintained on their corresponding RNAi treatments until the end of reproduction. We replicated this experiment by maintaining all experimental worms on empty vector RNAi producing *E. coli* during late reproduction (days 8-14). This was done to control for any possible effect that *daf-2* RNAi might have on male fertility and reproductive behaviour (*37*). In order to control for these effects, we report the results from the second replicate in the main text. The results from the first replicate are included in the Supplementary Material.

For intergenerational effects, we paired two males with a single hermaphrodite on day 8 for 24 hours (P1 generation). We transferred the P1 worms to new plates and let them lay eggs for 24 hours. We collected 50 hermaphrodite offspring (F1 generation) per treatment from these plates and maintained them on ad libitum conditions by transferring to new plates every 24 hours. For reproduction, we allowed the eggs to hatch and develop for two days, killed the hatched larvae at 42°C for 3.5 hours and stored them for counting later. For lifespan, we monitored survival until the death of each worm and censored the ones lost during the experiment.

### Visualisation of DAF-16 nuclear localisation

We used the zIs356 [daf-16p::daf-16a/b::GFP + rol-6(su1006)] strain to observe DAF-16 localisation. Synchronised worms expressing DAF-16::GFP were maintained under four different treatments (control, IF, rIIS, and combined treatments; see Fig. 1) until days 1 and 7 when visualisation took place. For IF and combined treatments, visualization occurred following a 9-hour fasting period. Worms were anesthetized using 25 mM levamisole on a 2% agarose pad, with approximately 20-25 worms per slide. Data on DAF-16 localisation were collected as binomial outcomes. We classified worms exhibiting predominant nuclear DAF-16 localisation as ‘1’, and those with primarily cytoplasmic localisation as ‘0’. This binary categorization was chosen to quantitatively assess the influence of different treatments on DAF-16 translocation dynamics.

### RNA extraction

For gene expression analysis, we collected worms from four different treatments on days 1, 7 and 13. On days 1 and 7, IF and rIIS + IF worms were collected right after the 9-hour fasting period, and the Control and rIIS worms were collected around the same time for consistency, although they were not fasted. We had three biological replicates with a sample size of 100. In total, we collected 1200 worms (4 treatments x 3 timepoints x 3 biological replicates). RNA extraction was performed using TRIsure (Bioline, Cat. No. BIO-38032) following standard phenol-chloroform RNA extraction protocol. The purity of the isolated RNA was checked with Nanodrop, and the integrity was quantified using Agilent 2200 TapeStation System.

RNA concentration was determined with Qubit using RNA HS Assay (Invitrogen™ Q33224).

### Illumina RNA Sequencing

500 ng of total RNA was used to perform mRNA isolation using NEBNext Poly(A) mRNA Magnetic Isolation Module (NEB Cat. No. E7490). Isolated mRNA material was used to prepare the libraries using NEBNext® Ultra II Directional RNA Library Prep Kit for Illumina® (NEB, Cat No. E7760S) following the manufacturer’s instructions. The resulting libraries were quantified using Qubit dsDNA HS Assay (Invitrogen™ Q32851).

### Data processing

Standard adapter sequences, common contaminants were removed from the paired-end reads using BBduk from the BBtools package version 37.62 (*55*). Positions with a Phred quality score of less than 15 were trimmed, and only reads with a length greater than 34 were retained. The quality of the reads was assessed using FastQC version v0.12.1 and MultiQC version 1.13 (*56*, *57*). The reads were aligned to the WBcel235 Caernohabditis elegans reference genome using STAR version 2.7.8a (*58*) with default parameters, and a splice junction database was generated from the Ensenbl release 107 reference annotation. Gene-level read counts were quantified using feature Counts version v2.0.3 (*59*) in paired-end mode.

### Differential expression analysis

Differential gene expression analysis was performed using DESeq2 version 1.32.0 (*60*). Pairwise comparisons were conducted using a Wald test, comparing different combinations of genotypes (empty vector or *daf-2* RNAi), diets (ad libitum or intermittent fasting), and time points (days 1, 7, and 13) against each other. The independent filtering feature was used to eliminate weakly expressed genes. P-values were corrected for multiple testing using the false discovery rate (FDR) computed using the Benjamini-Hochberg method. Log fold change shrinkage with the “ashr” shrinkage estimator (*61*) was utilized to control the log2 fold change estimates of genes with little information or high dispersion. Differentially expressed genes were identified based on log2 fold change cutoffs of 1 and FDR cutoff of 0.05.

### Gene Ontology analysis

Gene Ontology enrichment analysis was performed separately for uniquely and commonly up- and down-regulated genes using clusterProfiler version 4.2.0 (*62*) together with org.Ce.eg.db version 3.17.0 (*63*). P-values were corrected for multiple testing using the false discovery rate (FDR) computed using the Benjamini-Hochberg method. A 0.1 (10%) FDR threshold was applied to identify enriched biological processes relative to the background. Biological processes were deemed redundant if their GO terms consisted of overlapping sets of genes and belonged to the same branch. To avoid redundancy, only the top parent terms (superset of child term) were preserved, and all child terms (subset of parent term) were filtered out from the results.

### Statistical analysis

Self-fertilised reproduction - We fit generalized linear mixed-effects models using the glmmTMB package in R, with reproduction as the response variable, and food (AL vs. IF), RNAi (empty vector RNAi vs. rIIS) and their interaction as fixed effects. We also included batch as a fixed effect and the worm id as a random effect to account for any potential batch and worm id effects. We first used different families of models including Poisson, negative binomial type 1 and 2 and generalized Poisson distribution. Then, we compared the models using Akaike information criterion (AIC) and selected the best model based on the lowest AIC score. Finally, we used ANOVA on the best model to further investigate the significance of the fixed effects. This was done for all three populations (N2, USA and Portugal) separately.

Late-life mated reproduction – We fit generalized linear mixed-effects models using the glmmTMB package in R, with reproduction as the response variable, and food, RNAi and their interaction as fixed effects. We compared models with error distributions from four different families of models (Poisson, negative binomial type 1 and 2 and generalized Poisson distribution) and additional zero-inflation parameters. We selected the best model based on lowest AIC and performed ANOVA on it.

DAF-16 localisation – We used a generalized linear mixed-effects model with a binomial family to analyse the relationship between DAF-16 nuclear localisation and the interaction between RNAi, food, and day, while controlling for the worm id. We used ANOVA to test for the significance of fixed effects in our glmmTMB model.

We used the ‘glmmTMB’ package to fit generalized linear mixed-effects models using the ‘glmmTMB’ function, ‘car’ package to perform ANOVA analysis, ‘ggplot2’ package for data visualization, and ‘MuMIn’ package for model selection and inference (*64–67*). All analyses were performed in R v. 3.6.2 (*68*)

## Acknowledgements

Some strains were provided by the CGC, which is funded by NIH Office of Research Infrastructure Programs (P40 OD010440). We thank Wormbase for providing access to the *C. elegans* genome, annotation and genetic resources.

The research presented in this paper was carried out on the High Performance Computing Cluster supported by the Research and Specialist Computing Support service at the University of East Anglia.

## Funding

ZS was supported by Leverhulme Trust Early Career Fellowship.

AAM was supported by ERC Consolidator Grant (GermlineAgeingSoma 724909) and NERC NE/W001020/1.

AA and KH were supported by a UKRI Future Leaders Fellowship awarded to AA (grant number MR/S033769/1).

AS was supported by the UKRI Biotechnology and Biological Sciences Research Council Norwich Research Park Biosciences Doctoral Training Partnership (grant number BB/T008717/).

### Author Contributions

Conceptualization: ZS, AS, KH, AA and AAM

Methodology: ZS, KH, HC, ZC and DC

Data Analysis: ZS and AS

Writing - original draft: ZS and AAM

Writing - review & editing: ZS, AAM, AA, AS, KH, HC, ZC and DC.

### Competing interests

Authors declare that they have no competing interests.

### Data and materials availability

All data is shown in the main text or the supplementary materials. Raw data will be available on Dryad.

---

**Fig. S1.**
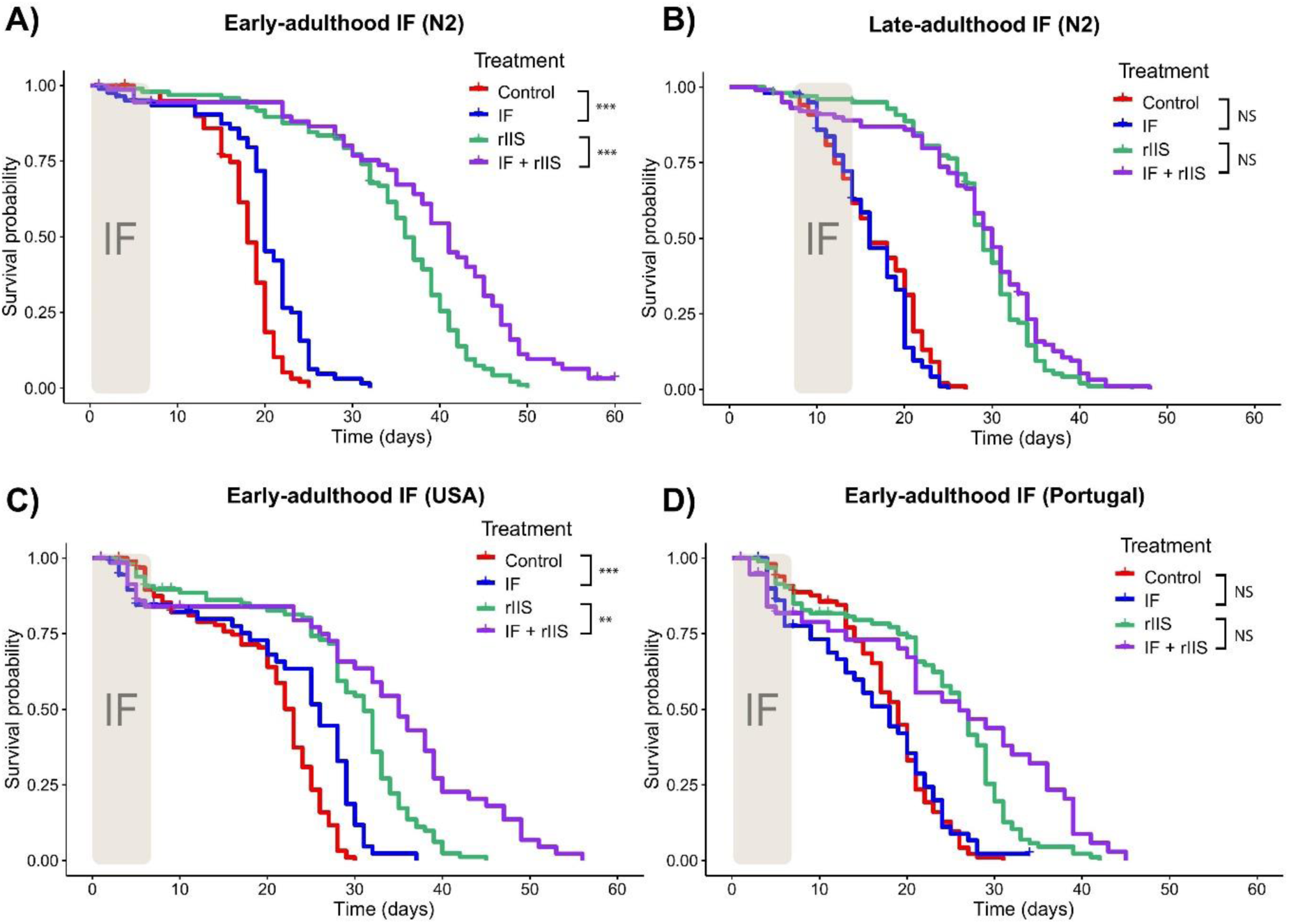
Effect of IF and rIIS on survival including matricides. Effect of early and late-adulthood IF on survival of N2 worms with and without rIIS (A and B, respectively). Effect of early-adulthood IF on survival of USA and Portugal strains with and without rIIS (C and D, respectively). Significance in the divergence of survival curves was assessed using log-rank tests, with statistical significance represented as follows: NS indicates P > 0.05, * indicates P ≤ 0.05, ** indicates P ≤ 0.01, and *** indicates P ≤ 0.001.

**Fig. S2.**
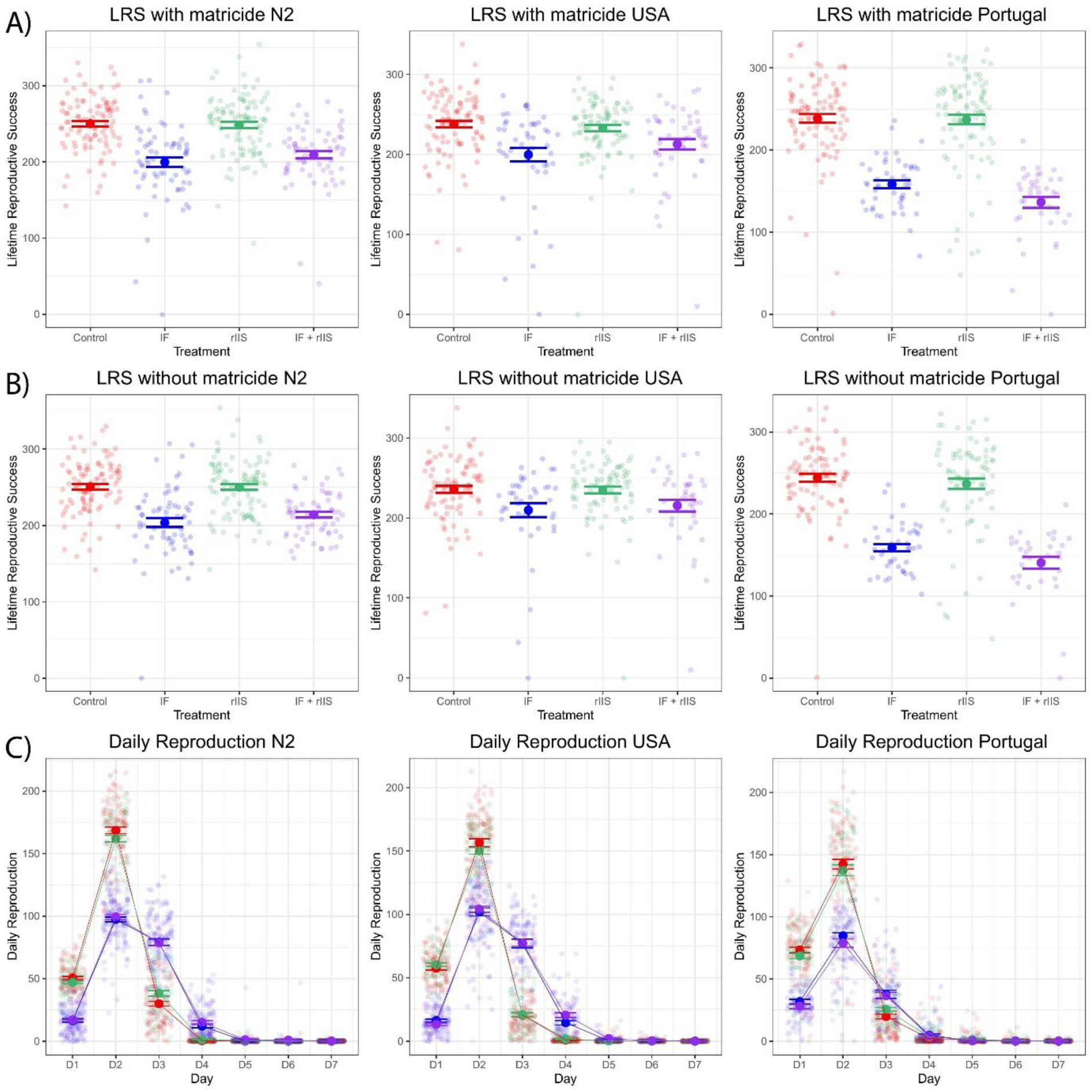
Effect of IF and rIIS on early-life reproduction. Early-life reproduction including matricie across three different populations (A), early-life reproduction without matricide across three different populations (B), daily reproduction across three different populations (C).

**Fig. S3.**
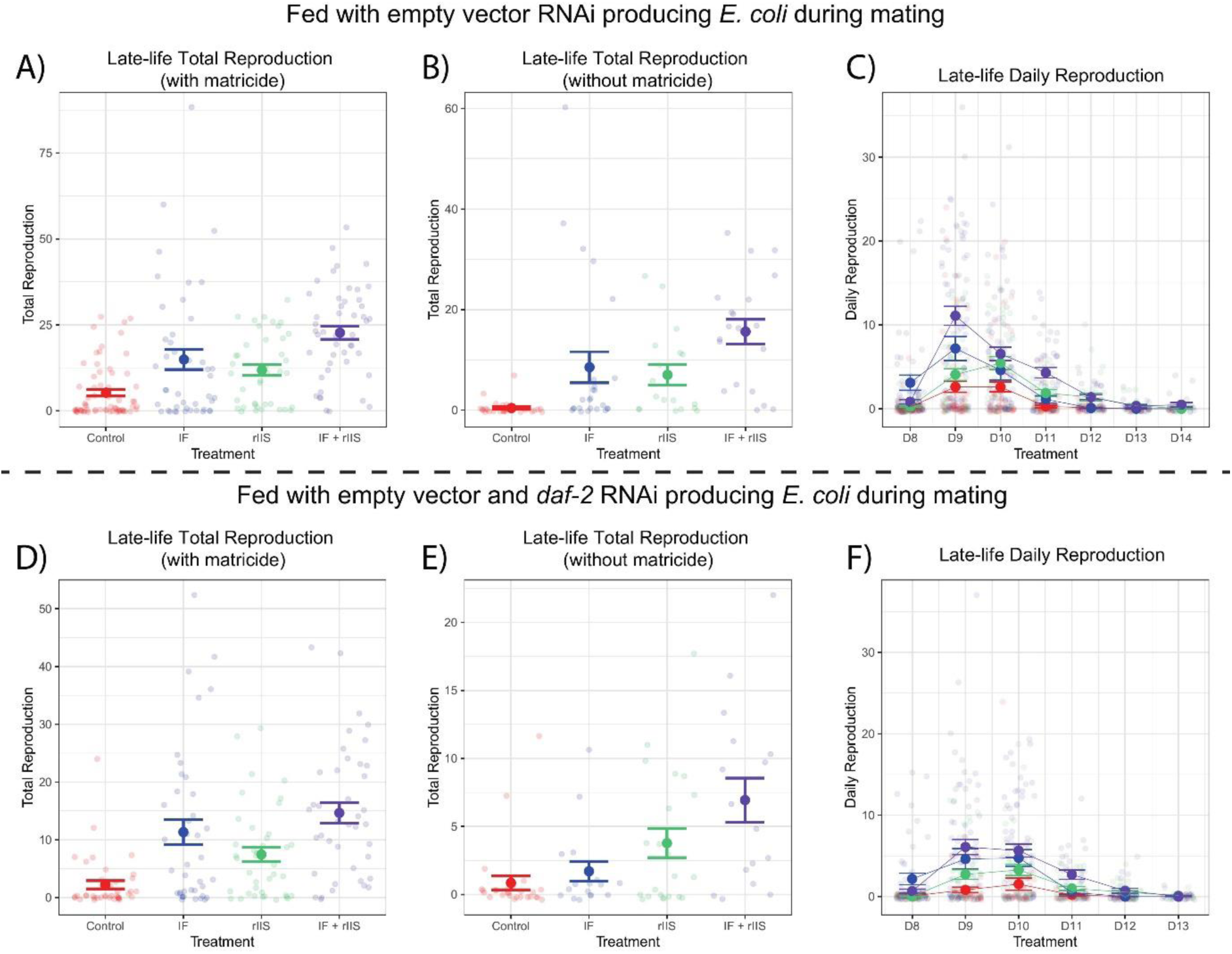
Effect of IF and rIIS on late-life reproduction. Late-life reproduction including matricide (A), late-life reproduction without matricide (B), daily reproduction (C) in worms fed with empty vector RNAi producing *E. coli* during mating. Late-life reproduction including matricide (D), late-life reproduction without matricide (E), daily reproduction (F) in worms fed with empty vector and *daf-2* RNAi producing *E. coli* during mating.

**Fig. S4.**
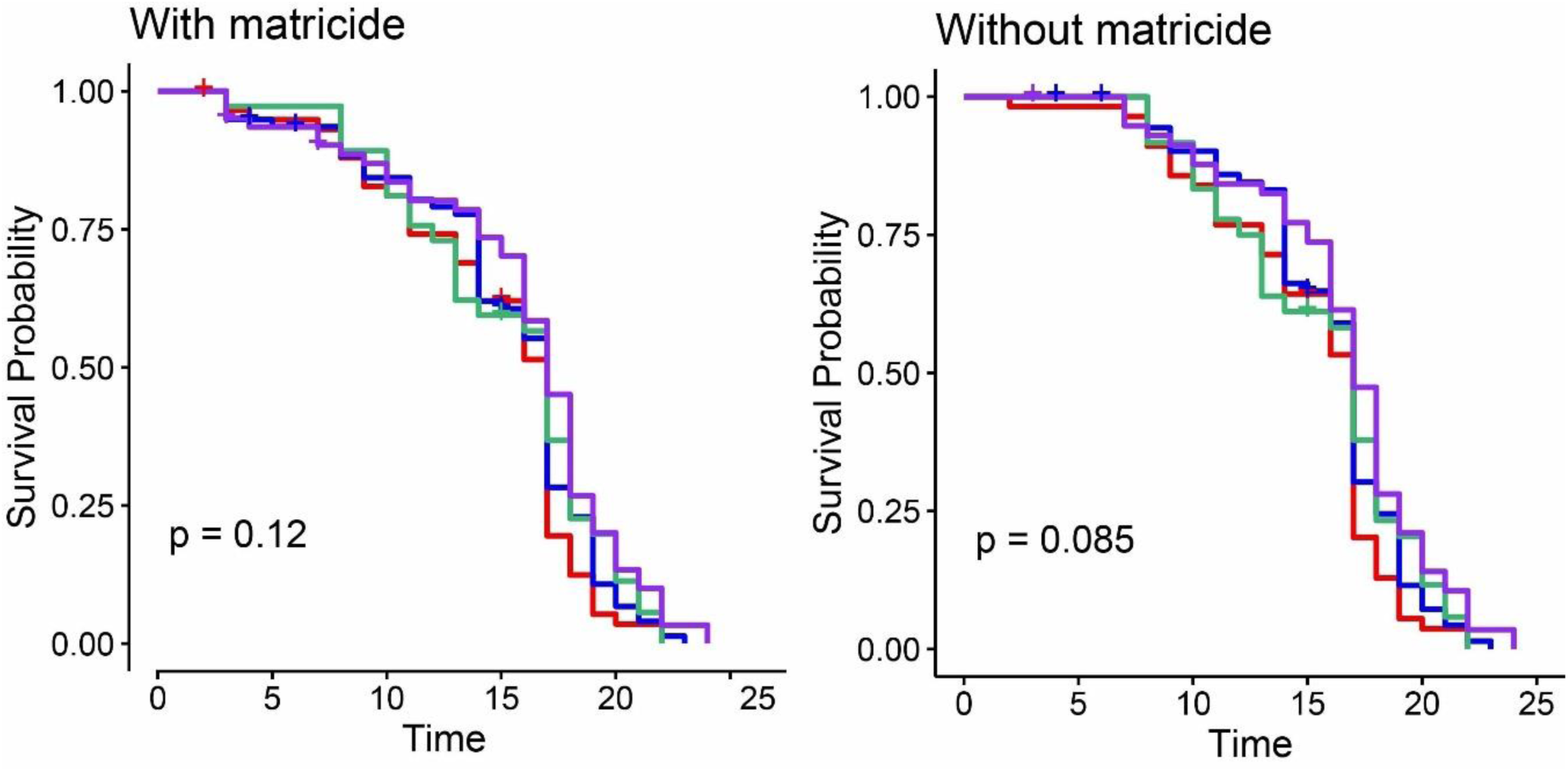
Effect of IF and rIIS on offspring survival. Survival of offspring from parents with Control, IF, rIIS and IF + rIIS treatments. Survival with matricide (A) and without matricide (B). Significance in the divergence of survival curves was assessed using log-rank tests, with statistical significance represented as follows: NS indicates P > 0.05, * indicates P ≤ 0.05, ** indicates P ≤ 0.01, and *** indicates P ≤ 0.001.

**Fig. S5.**
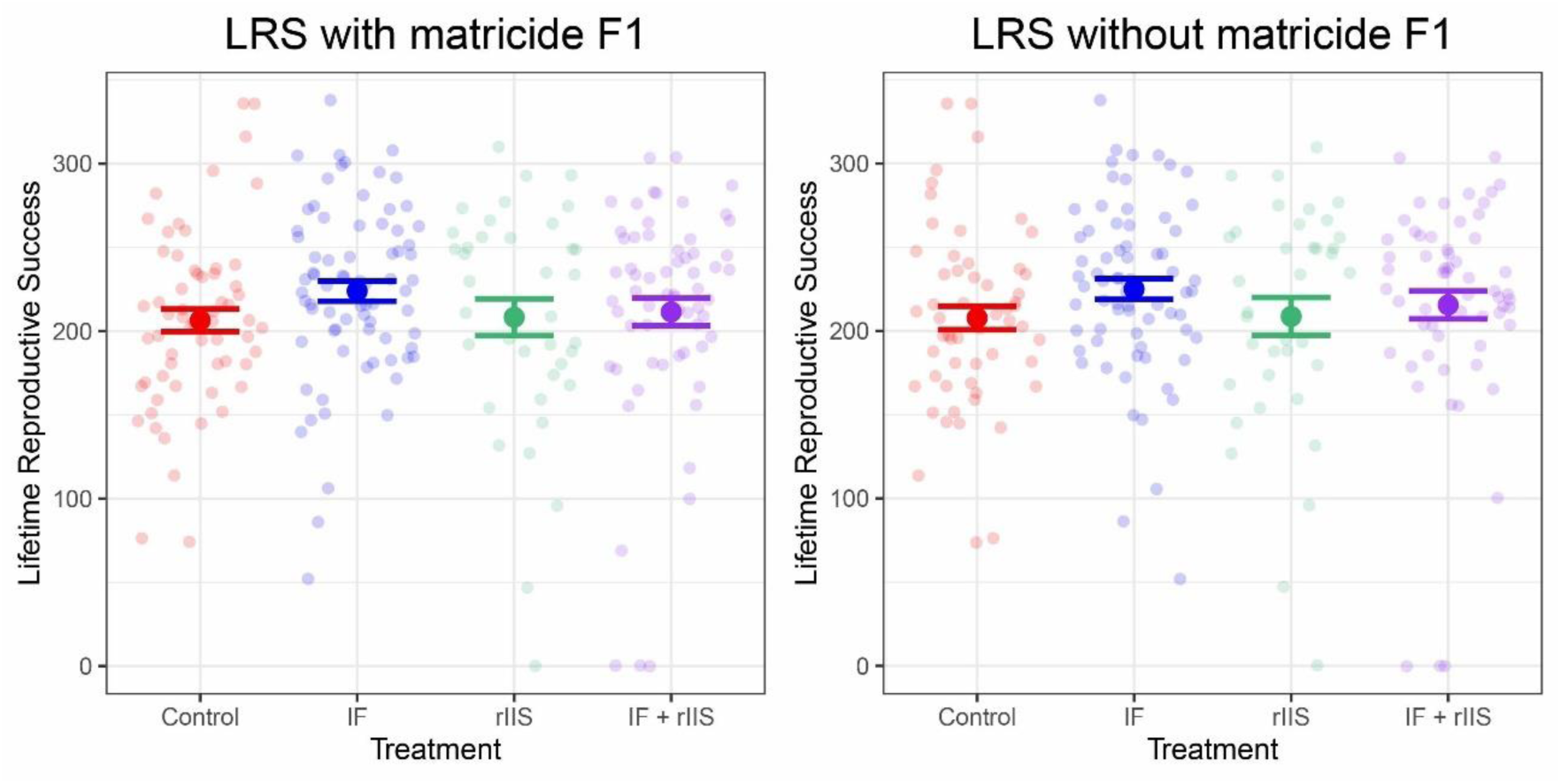
Effect of IF and rIIS on offspring lifetime reproductive success. Lifetime reproductive success of offspring from parents with Control, IF, rIIS and IF + rIIS treatments. LRS with matricide (A), LRS without matricide (B).

**Fig. S6.**
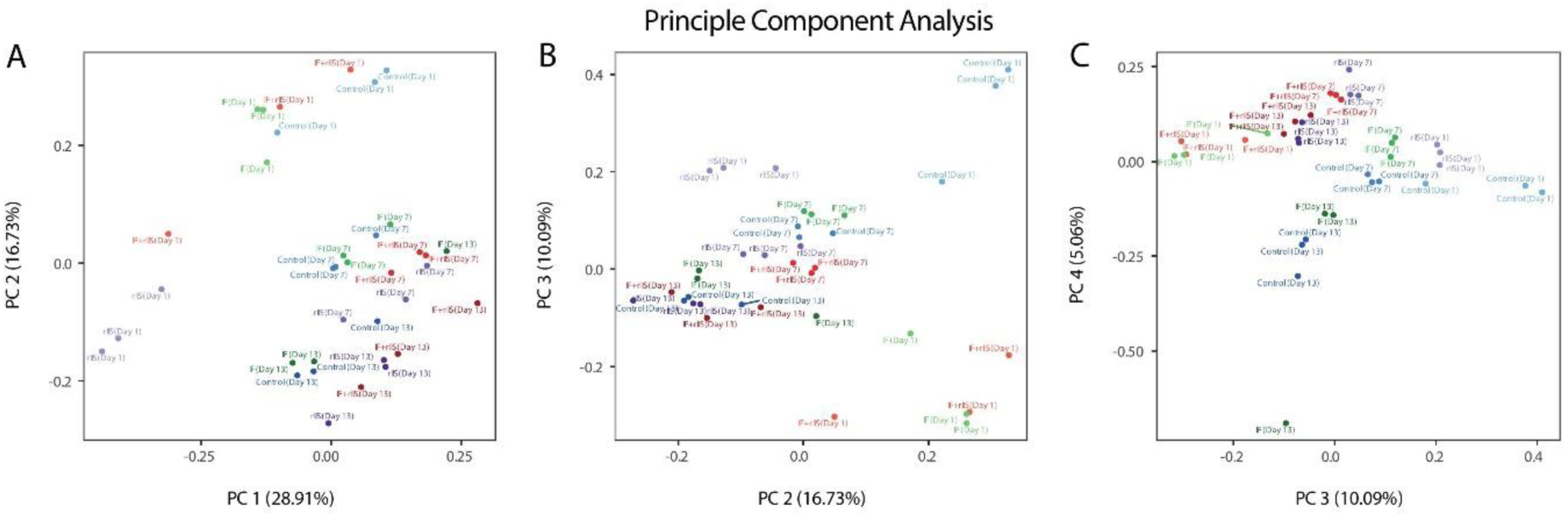
Principal Component Analysis (PCA) of RNA-seq Samples. PCA plots illustrating the gene expression profiles for samples under four different treatments: Control, IF, rIIS, and Combined (IF + rIIS). The plots compare the gene expression patterns at three distinct time points – days 1, 7, and 13 – to demonstrate how the treatments influence genetic activity over time.

**Fig. S7.**
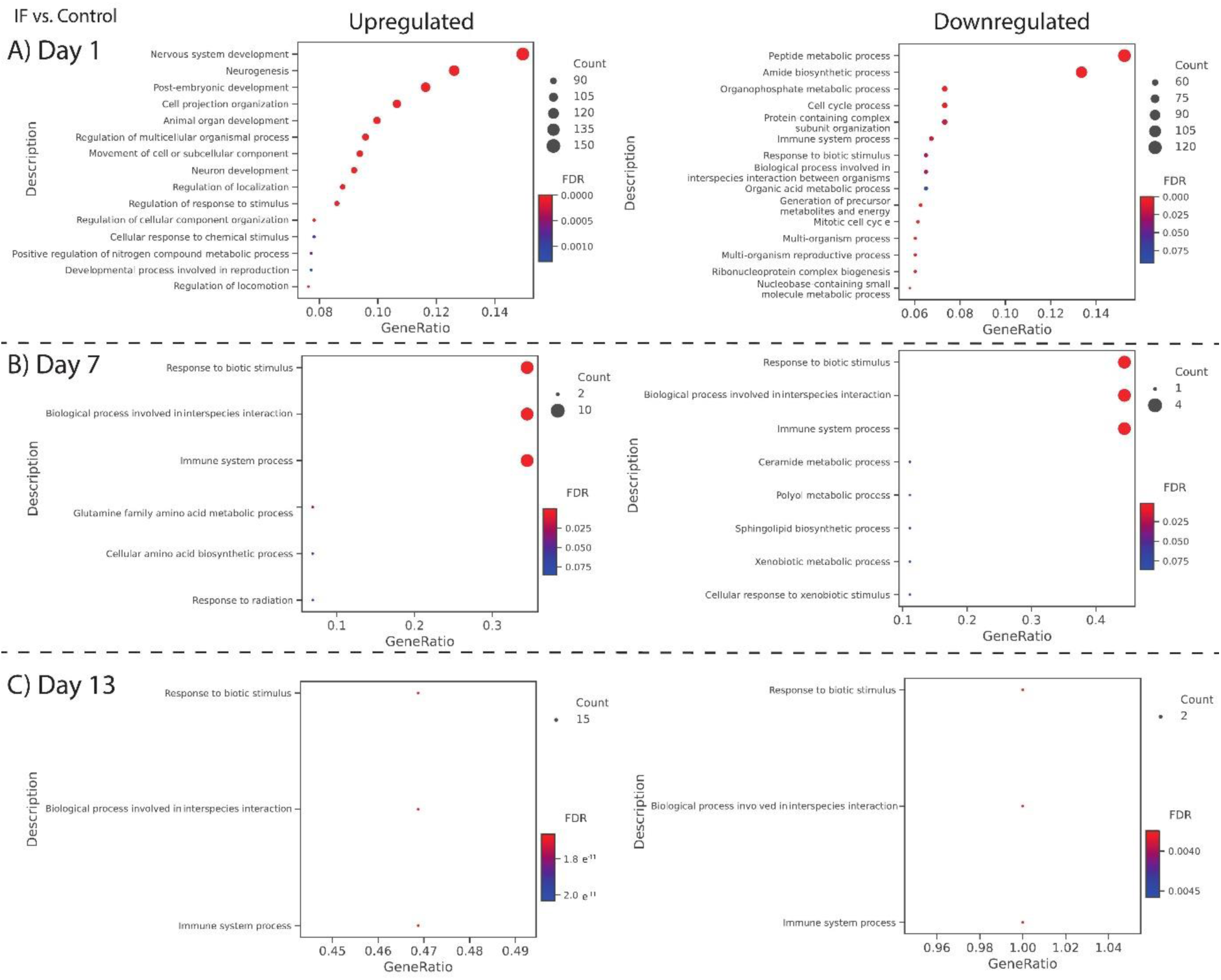
Gene Ontology analysis of differentially expressed genes in the IF treatment. GO term analyses showing the functions of genes up and down-regulated by the IF treatment compared to control across day 1 (A), day 7 (B) and day 13 (C).

**Fig. S8.**
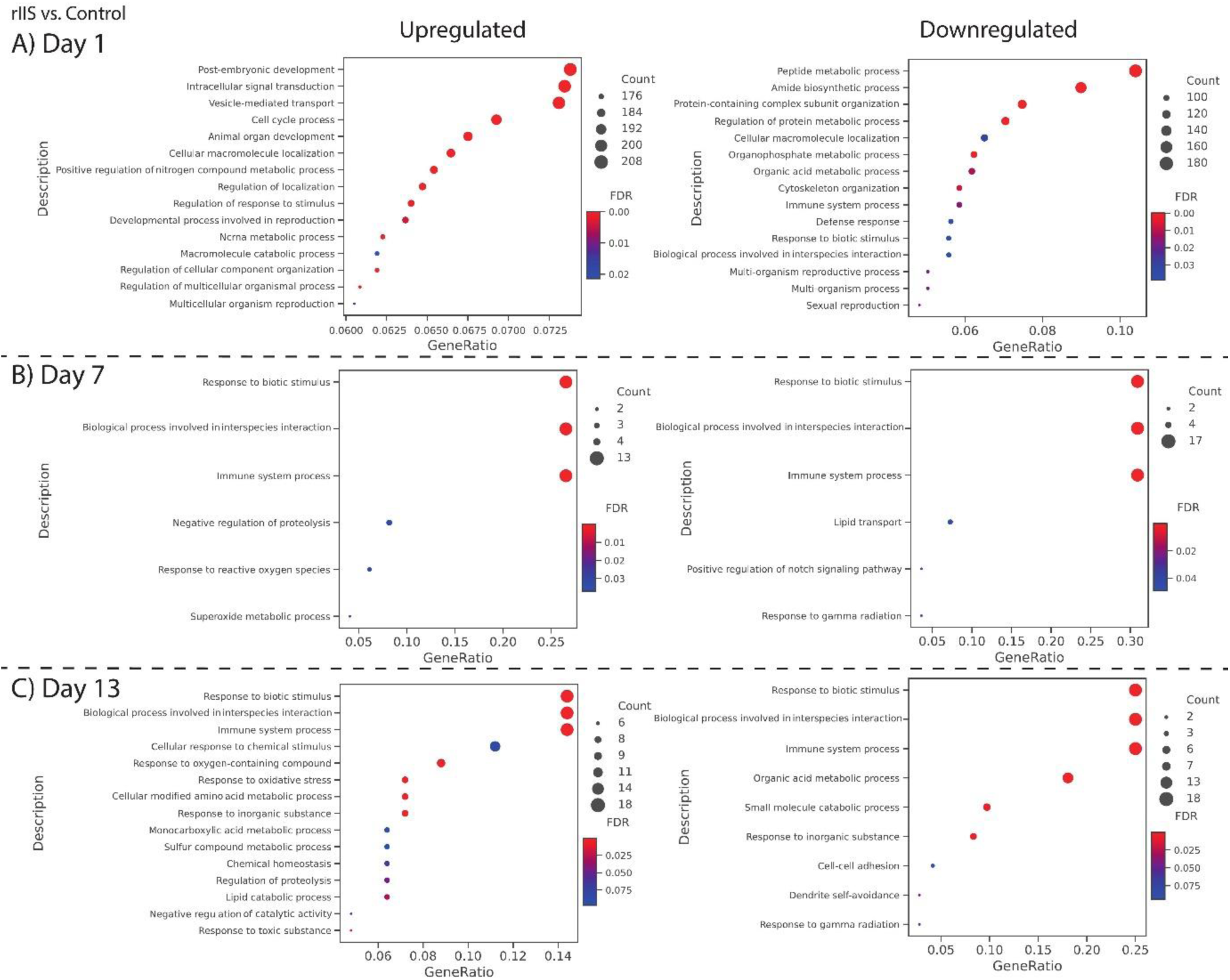
Gene Ontology analysis of differentially expressed genes in the rIIS treatment. GO term analyses showing the functions of genes up and down-regulated by the rIIS treatment compared to control across day 1 (A), day 7 (B) and day 13 (C).

## Notes

### Competing Interest Statement

The authors have declared no competing interest.

### Summary of Updates

We included new data on the intergenerational effects of rIIS and IF on offspring reproduction and survival. With the addition of these new results, we made additions and slight changes to the relevant sections in a way to underline the importance of our findings with respect to the evolutionary theories of ageing.

## References

1. W. D. Hamilton, The moulding of senescence by natural selection. J. Theor. Biol. 12, 12–45 (1966).

2. P. B. Medawar, An unsolved problem of biology. (1952).

3. G. C. Williams, Pleiotropy, Natural Selection, and the Evolution of Senescence. Evolution 11, 398–411 (1957).

4. J. F. Lemaître, J. Moorad, J.-M. Gaillard, A. A. Maklakov, D. H. Nussey, A unified framework for evolutionary genetic and physiological theories of aging. PLOS Biol. 22, e3002513 (2024).

5. T. B. L. Kirkwood, Evolution of ageing. Nature 270, 301–304 (1977).

6. T. B. L. Kirkwood, R. Holliday, The evolution of ageing and longevity. Proc. R. Soc. Lond. B Biol. Sci. 205, 531–546 (1979).

7. T. B. L. Kirkwood, M. R. Rose, Evolution of senescence: late survival sacrificed for reproduction. Philos. Trans. R. Soc. Lond. B. Biol. Sci. 332, 15–24 (1991).

8. J. P. de Magalhães, Programmatic features of aging originating in development: aging mechanisms beyond molecular damage? FASEB J. 26, 4821–4826 (2012).

9. J. P. de Magalhães, G. M. Church, Genomes Optimize Reproduction: Aging as a Consequence of the Developmental Program. Physiology 20, 252–259 (2005).

10. D. Gems, The hyperfunction theory: An emerging paradigm for the biology of aging. Ageing Res. Rev. 74, 101557 (2022).

11. A. A. Maklakov, T. Chapman, Evolution of ageing as a tangle of trade-offs: energy versus function. Proc. R. Soc. B Biol. Sci. 286, 20191604 (2019).

12. S. Luo, G. A. Kleemann, J. M. Ashraf, W. M. Shaw, C. T. Murphy, TGF-β and Insulin Signaling Regulate Reproductive Aging via Oocyte and Germline Quality Maintenance. Cell 143, 299–312 (2010).

13. S. Luo, W. M. Shaw, J. Ashraf, C. T. Murphy, TGF-ß Sma/Mab signaling mutations uncouple reproductive aging from somatic aging. PLoS Genet. 5, e1000789 (2009).

14. C. Huang, C. Xiong, K. Kornfeld, Measurements of age-related changes of physiological processes that predict lifespan of Caenorhabditis elegans. Proc. Natl. Acad. Sci. 101, 8084–8089 (2004).

15. S. E. Hughes, K. Evason, C. Xiong, K. Kornfeld, Genetic and Pharmacological Factors That Influence Reproductive Aging in Nematodes. PLoS Genet. 3, e25 (2007).

16. C. Li, H. Zhang, H. Wu, R. Li, D. Wen, Y. Tang, Z. Gao, R. Xu, S. Lu, Q. Wei, X. Zhao, M. Pan, B. Ma, Intermittent fasting reverses the declining quality of aged oocytes. Free Radic. Biol. Med. 195, 74–88 (2023).

17. Z. Sultanova, E. R. Ivimey-Cook, T. Chapman, A. A. Maklakov, Fitness benefits of dietary restriction. Proc. R. Soc. B Biol. Sci. 288, 20211787 (2021).

18. E. L. Greer, A. Brunet, Different dietary restriction regimens extend lifespan by both independent and overlapping genetic pathways in C. elegans. Aging Cell 8, 113–127 (2009).

19. S. T. Henderson, T. E. Johnson, daf-16 integrates developmental and environmental inputs to mediate aging in the nematode Caenorhabditis elegans. Curr. Biol. 11, 1975– 1980 (2001).

20. M. P. Mattson, V. D. Longo, M. Harvie, Impact of intermittent fasting on health and disease processes. Ageing Res. Rev. 39, 46–58 (2017).

21. C. Baumeier, D. Kaiser, J. Heeren, L. Scheja, C. John, C. Weise, M. Eravci, M. Lagerpusch, G. Schulze, H.-G. Joost, R. W. Schwenk, A. Schürmann, Caloric restriction and intermittent fasting alter hepatic lipid droplet proteome and diacylglycerol species and prevent diabetes in NZO mice. Biochim. Biophys. Acta BBA - Mol. Cell Biol. Lipids 1851, 566–576 (2015).

22. J. H. Catterson, M. Khericha, M. C. Dyson, A. J. Vincent, R. Callard, S. M. Haveron, A. Rajasingam, M. Ahmad, L. Partridge, Short-Term, Intermittent Fasting Induces Long-Lasting Gut Health and TOR-Independent Lifespan Extension. Curr. Biol. 28, 1714– 1724 (2018).

23. S. Honjoh, T. Yamamoto, M. Uno, E. Nishida, Signalling through RHEB-1 mediates intermittent fasting-induced longevity in C. elegans. Nature 457, 726–730 (2009).

24. B. Liu, A. J. Page, G. Hatzinikolas, M. Chen, G. A. Wittert, L. K. Heilbronn, Intermittent Fasting Improves Glucose Tolerance and Promotes Adipose Tissue Remodeling in Male Mice Fed a High-Fat Diet. Endocrinology 160, 169–180 (2019).

25. M. Ulgherait, A. M. Midoun, S. J. Park, J. A. Gatto, S. J. Tener, J. Siewert, N. Klickstein, J. C. Canman, W. W. Ja, M. Shirasu-Hiza, Circadian autophagy drives iTRF-mediated longevity. Nature 598, 353–358 (2021).

26. M. Uno, S. Honjoh, M. Matsuda, H. Hoshikawa, S. Kishimoto, T. Yamamoto, M. Ebisuya, T. Yamamoto, K. Matsumoto, E. Nishida, A Fasting-Responsive Signaling Pathway that Extends Life Span in C. elegans. Cell Rep. 3, 79–91 (2013).

27. D. Weinkove, J. R. Halstead, D. Gems, N. Divecha, Long-term starvation and ageing induce AGE-1/PI 3-kinase-dependent translocation of DAF-16/FOXO to the cytoplasm. BMC Biol. 4, 1–13 (2006).

28. X. Sun, W. D. Chen, Y. D. Wang, DAF-16/FOXO transcription factor in aging and longevity. Front. Pharmacol. 8, 548 (2017).

29. E. M. L. Duxbury, H. Carlsson, K. Sales, Z. Sultanova, S. Immler, T. Chapman, A. A. Maklakov, Multigenerational downregulation of insulin/IGF-1 signaling in adulthood improves lineage survival, reproduction, and fitness in Caenorhabditis elegans supporting the developmental theory of ageing. Evolution 76, 2829–2845 (2022).

30. E. Schuster, J. J. McElwee, J. M. A. Tullet, R. Doonan, F. Matthijssens, J. S. Reece-Hoyes, I. A. Hope, J. R. Vanfleteren, J. M. Thornton, D. Gems, DamID in C. elegans reveals longevity-associated targets of DAF-16/FoxO. Mol. Syst. Biol. 6, 399 (2010).

31. X. Huang, W. Pan, W. Kim, A. White, S. Li, H. Li, K. Lee, B. B. Fuchs, K. Zeng, E. Mylonakis, Caenorhabditis elegans mounts a p38 MAPK pathway-mediated defence to Cutibacterium acnes infection. Cell. Microbiol. 22, e13234 (2020).

32. X. Zhang, Y. Zhang, Neural-Immune Communication in Caenorhabditis elegans. Cell Host Microbe 5, 425–429 (2009).

33. Y. Weng, S. Zhou, K. Morillo, R. Kaletsky, S. Lin, C. M.- BioRxiv, U. 2023, Analysis of the Neuron-specific IIS/FOXO Transcriptome in Aged Animals Reveals Regulators of Neuronal and Cognitive Aging. *biorxiv*.org 2023–07 (2023).

34. S. Alper, S. J. McBride, B. Lackford, J. H. Freedman, D. A. Schwartz, Specificity and Complexity of the *Caenorhabditis elegans* Innate Immune Response. Mol. Cell. Biol. 27, 5544–5553 (2007).

35. H. A. Tissenbaum, G. Ruvkun, An Insulin-like Signaling Pathway Affects Both Longevity and Reproduction in Caenorhabditis elegans. Genetics 148, 703–717 (1998).

36. P. L. Larsen, P. S. Albert, D. L. Riddle, Genes that regulate both development and longevity in Caenorhabditis elegans. Genetics 139, 1567–1583 (1995).

37. D. Gems, A. J. Sutton, M. L. Sundermeyer, P. S. Albert, K. V. King, M. L. Edgley, P. L. Larsen, D. L. Riddle, Two pleiotropic classes of daf-2 mutation affect larval arrest, adult behavior, reproduction and longevity in Caenorhabditis elegans. Genetics 150, 129–155 (1998).

38. N. M. Templeman, S. Luo, R. Kaletsky, C. Shi, J. Ashraf, W. Keyes, C. T. Murphy, Insulin Signaling Regulates Oocyte Quality Maintenance with Age via Cathepsin B Activity. Curr. Biol. 28, 753–760 (2018).

39. J. Chen, E. E. Lewis, J. R. Carey, H. Caswell, E. P. Caswell-Chen, The ecology and biodemography of *Caenorhabditis elegans*. Exp. Gerontol. 41, 1059–1065 (2006).

40. A. R. Mendenhall, D. Wu, S. K. Park, J. R. Cypser, P. M. Tedesco, C. D. Link, P. C. Phillips, T. E. Johnson, Genetic dissection of late-life fertility in Caenorhabditis elegans. J. Gerontol. - Ser. Biol. Sci. Med. Sci. 66, 842–854 (2011).

41. M. I. Lind, S. Ravindran, Z. Sekajova, H. Carlsson, A. Hinas, A. A. Maklakov, Experimentally reduced insulin/IGF-1 signaling in adulthood extends lifespan of parents and improves Darwinian fitness of their offspring. Evol. Lett. 3, 207–216 (2019).

42. H. Carlsson, E. Ivimey-Cook, E. M. L. Duxbury, N. Edden, K. Sales, A. A. Maklakov, Ageing as “early-life inertia”: Disentangling life-history trade-offs along a lifetime of an individual. Evol. Lett. 5, 551–564 (2021).

43. M. Hansen, S. Taubert, D. Crawford, N. Libina, S. J. Lee, C. Kenyon, Lifespan extension by conditions that inhibit translation in Caenorhabditis elegans. Aging Cell 6, 95–110 (2007).

44. P. Syntichaki, K. Troulinaki, N. Tavernarakis, Protein synthesis is a novel determinant of aging in Caenorhabditis elegans. Ann. N. Y. Acad. Sci. 1119, 289–295 (2007).

45. M. L. Ladage, S. D. King, D. J. Burks, D. L. Quan, A. M. Garcia, R. K. Azad, P. A. Padilla, Glucose or Altered Ceramide Biosynthesis Mediate Oxygen Deprivation Sensitivity Through Novel Pathways Revealed by Transcriptome Analysis in Caenorhabditis elegans. G3 Genes Genomes Genet. 6, 3149–3160 (2016).

46. K. Z. Pan, J. E. Palter, A. N. Rogers, A. Olsen, D. Chen, G. J. Lithgow, P. Kapahi, Inhibition of mRNA translation extends lifespan in Caenorhabditis elegans. Aging Cell 6, 111–119 (2007).

47. T. Herman, E. Hartwieg, H. R. Horvitz, sqv mutants of Caenorhabditis elegans are defective in vulval epithelial invagination. Proc. Natl. Acad. Sci. 96, 968–973 (1999).

48. F. Licastro, G. Candore, D. Lio, E. Porcellini, G. Colonna-Romano, C. Franceschi, C. Caruso, Innate immunity and inflammation in ageing: A key for understanding age-related diseases. Immun. Ageing 2, 1–14 (2005).

49. P. Capilla-Lasheras, D. M. Dominoni, S. A. Babayan, P. J. O’Shaughnessy, M. Mladenova, L. Woodford, C. J. Pollock, T. Barr, F. Baldini, B. Helm, Elevated Immune Gene Expression Is Associated with Poor Reproductive Success of Urban Blue Tits. Front. Ecol. Evol. 5 (2017).

50. D. K. Fabian, K. Garschall, P. Klepsatel, G. Santos-Matos, É. Sucena, M. Kapun, B. Lemaitre, C. Schlötterer, R. Arking, T. Flatt, Evolution of longevity improves immunity in Drosophila. Evol. Lett. 2, 567–579 (2018).

51. C. Boehnisch, D. Wong, M. Habig, K. Isermann, N. K. Michiels, T. Roeder, R. C. May, H. Schulenburg, Protist-Type Lysozymes of the Nematode Caenorhabditis elegans Contribute to Resistance against Pathogenic Bacillus thuringiensis. PLOS ONE 6, e24619 (2011).

52. I. Gallotta, A. Sandhu, M. Peters, M. Haslbeck, R. Jung, S. Agilkaya, J. L. Blersch, C. Rödelsperger, W. Röseler, C. Huang, R. J. Sommer, D. C. David, Extracellular proteostasis prevents aggregation during pathogenic attack. Nature 584, 410–414 (2020).

53. E. Lionaki, N. Tavernarakis, High-Throughput and Longitudinal Analysis of Aging and Senescent Decline in Caenorhabditis elegans. Cell Senescence Methods Protoc., 485– 500 (2013).

54. A. Dillin, D. Crawford, C. Kenyon, U. 2002, Timing requirements for insulin/IGF-1 signaling in C. elegans. Science 298, 830–834 (2002).

55. B. Bushnell, BBMap. sourceforge. net/projects/bbmap (2020).

56. S. Andrews, FastQC: a quality control tool for high throughput sequence data (2010).

57. P. Ewels, M. Magnusson, S. Lundin, M. Käller, MultiQC: summarize analysis results for multiple tools and samples in a single report. Bioinformatics 32, 3047–3048 (2016).

58. A. Dobin, C. A. Davis, F. Schlesinger, J. Drenkow, C. Zaleski, S. Jha, P. Batut, M. Chaisson, T. R. Gingeras, STAR: ultrafast universal RNA-seq aligner. Bioinformatics 29, 15–21 (2013).

59. Y. Liao, G. K. Smyth, W. Shi, featureCounts: an efficient general purpose program for assigning sequence reads to genomic features. Bioinformatics 30, 923–930 (2014).

60. M. I. Love, W. Huber, S. Anders, Moderated estimation of fold change and dispersion for RNA-seq data with DESeq2. Genome Biol. 15, 550 (2014).

61. M. Stephens, False discovery rates: a new deal. Biostatistics 18, 275–294 (2017).

62. T. Wu, E. Hu, S. Xu, M. Chen, P. Guo, Z. Dai, T. Feng, L. Zhou, W. Tang, L. I. Zhan, clusterProfiler 4.0: A universal enrichment tool for interpreting omics data. The innovation 2 (2021).

63. M. Carlson, org. Mm. eg. db: Genome wide annotation for Mouse. R package version 3.2. 3. Bioconductor Lond. U. K. Genome Biol. BMC (2019).

64. K. Barton, M. K. Barton, Package ‘mumin.’ Version 1, 439 (2015).

65. J. Fox, S. Weisberg, D. Adler, D. Bates, G. Baud-Bovy, S. Ellison, D. Firth, M. Friendly, G. Gorjanc, S. Graves, Package ‘car.’ *Vienna R Found*. Stat. Comput. 16, 1–158 (2012).

66. A. Magnusson, H. J. Skaug, A. Nielsen, C. Berg, K. Kristensen, M. Maechler, K. van Bentham, B. M. Bolker, M. E. Brooks, Package ‘glmmTMB. R Package Version 01 (2019).

67. H. Wickham, ggplot2. *WIREs Comput*. Stat. 3, 180–185 (2011).

68. R Core Team, R version 3.6. 2: A language and environmental for statistical computing (2019).

