## Supplementary figures and images for "Optimising age-specific insulin signalling to slow down reproductive ageing increases fitness in different environments"

### Fig. S1

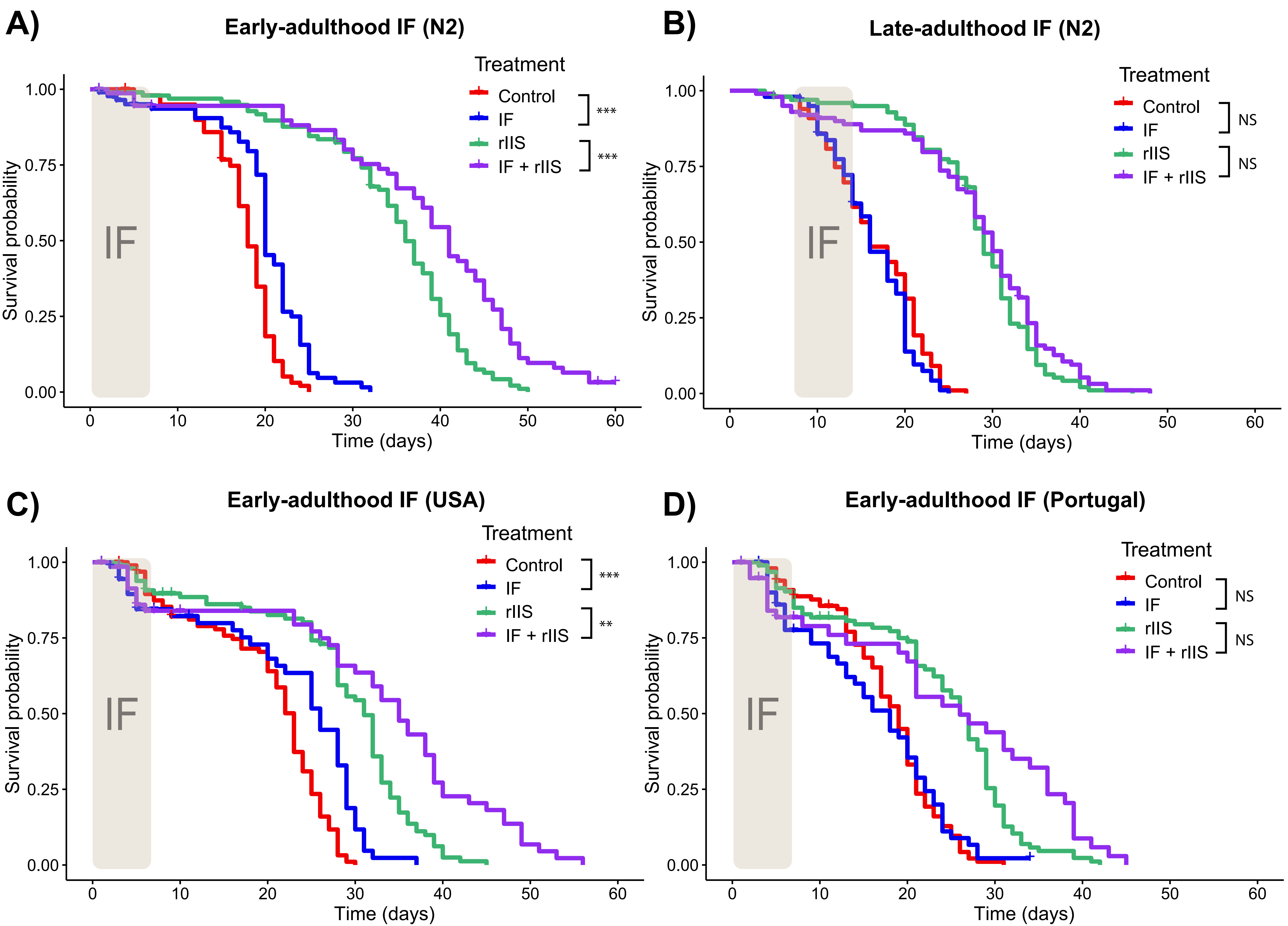

### Fig. S2

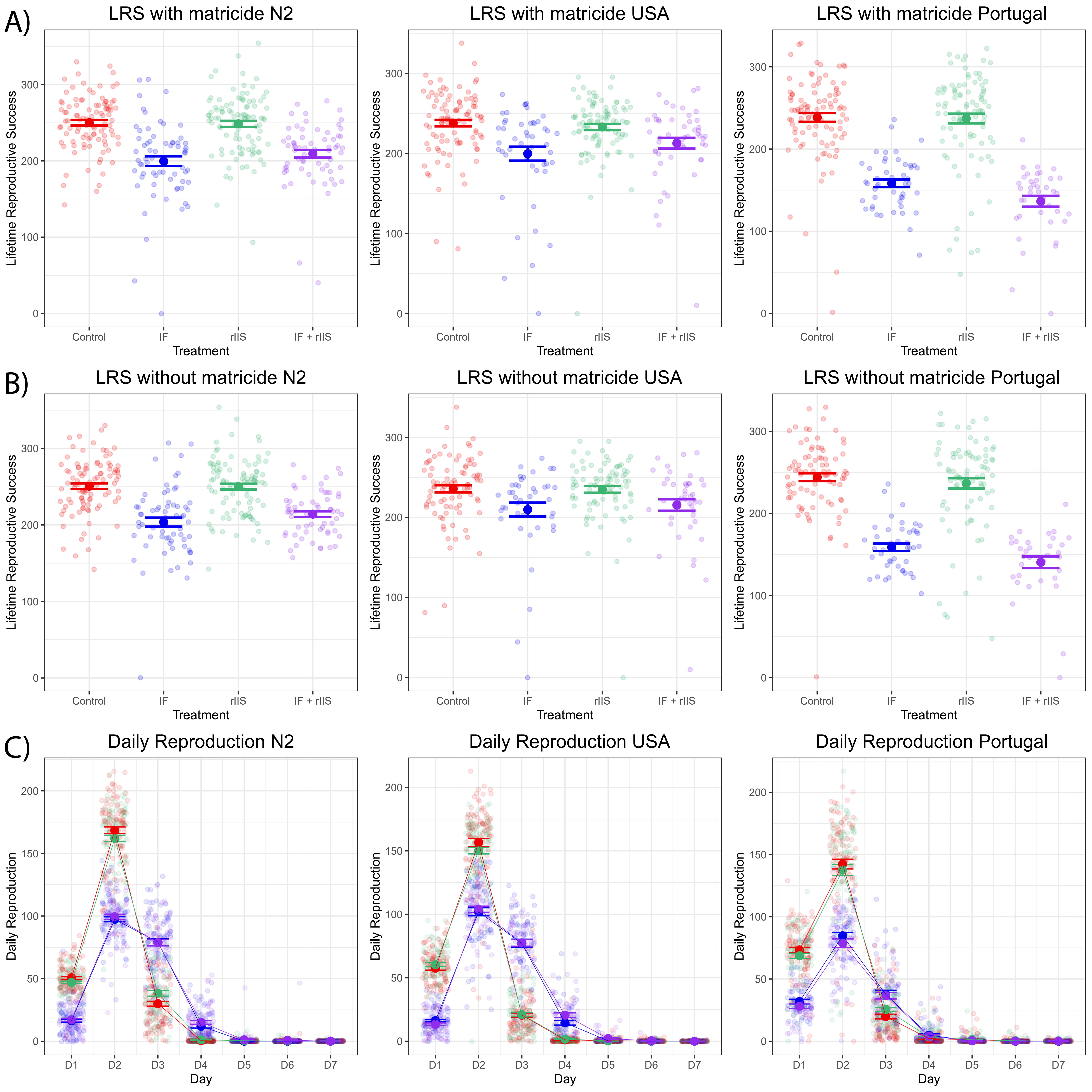

### Fig. S3

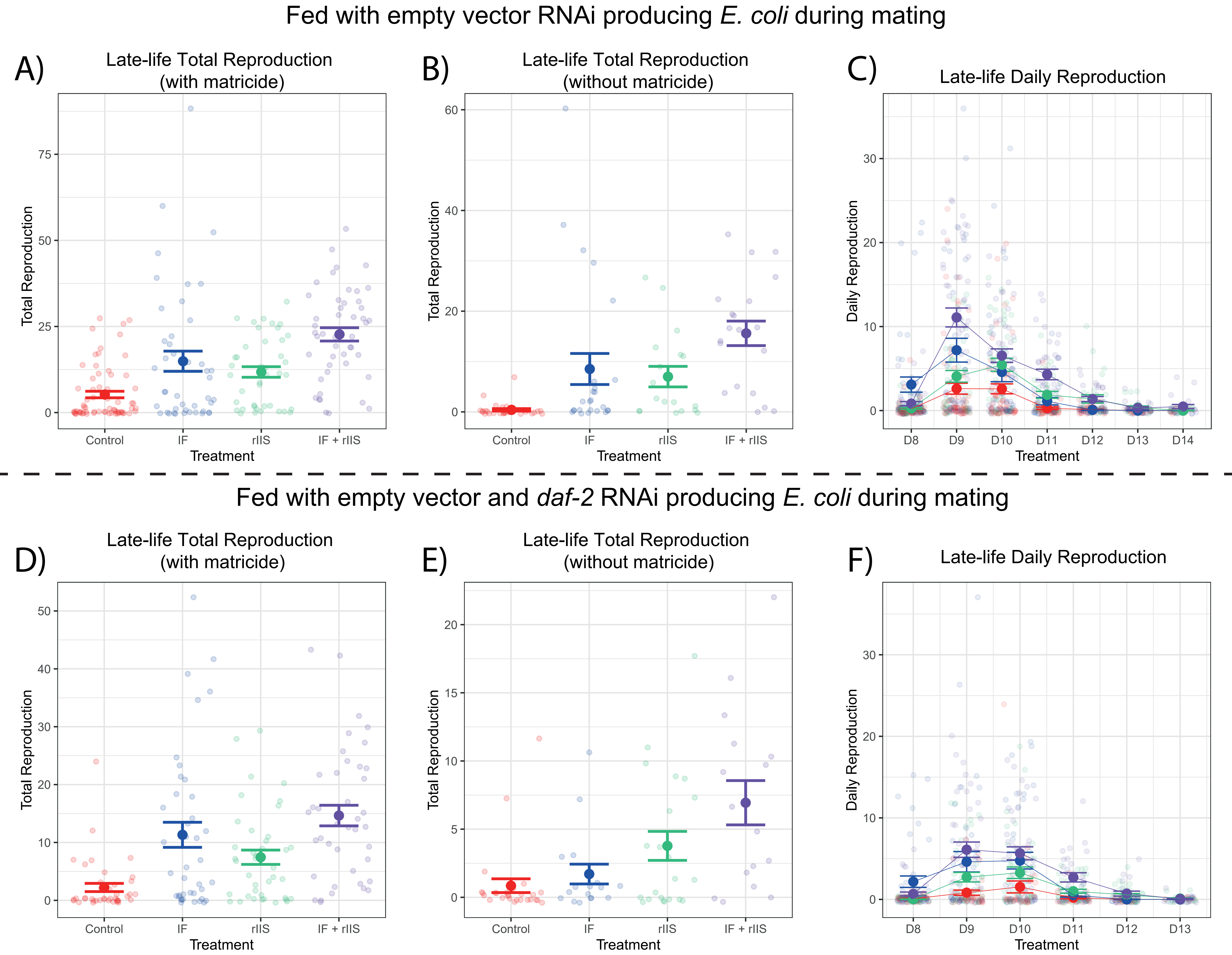

### Fig. S4

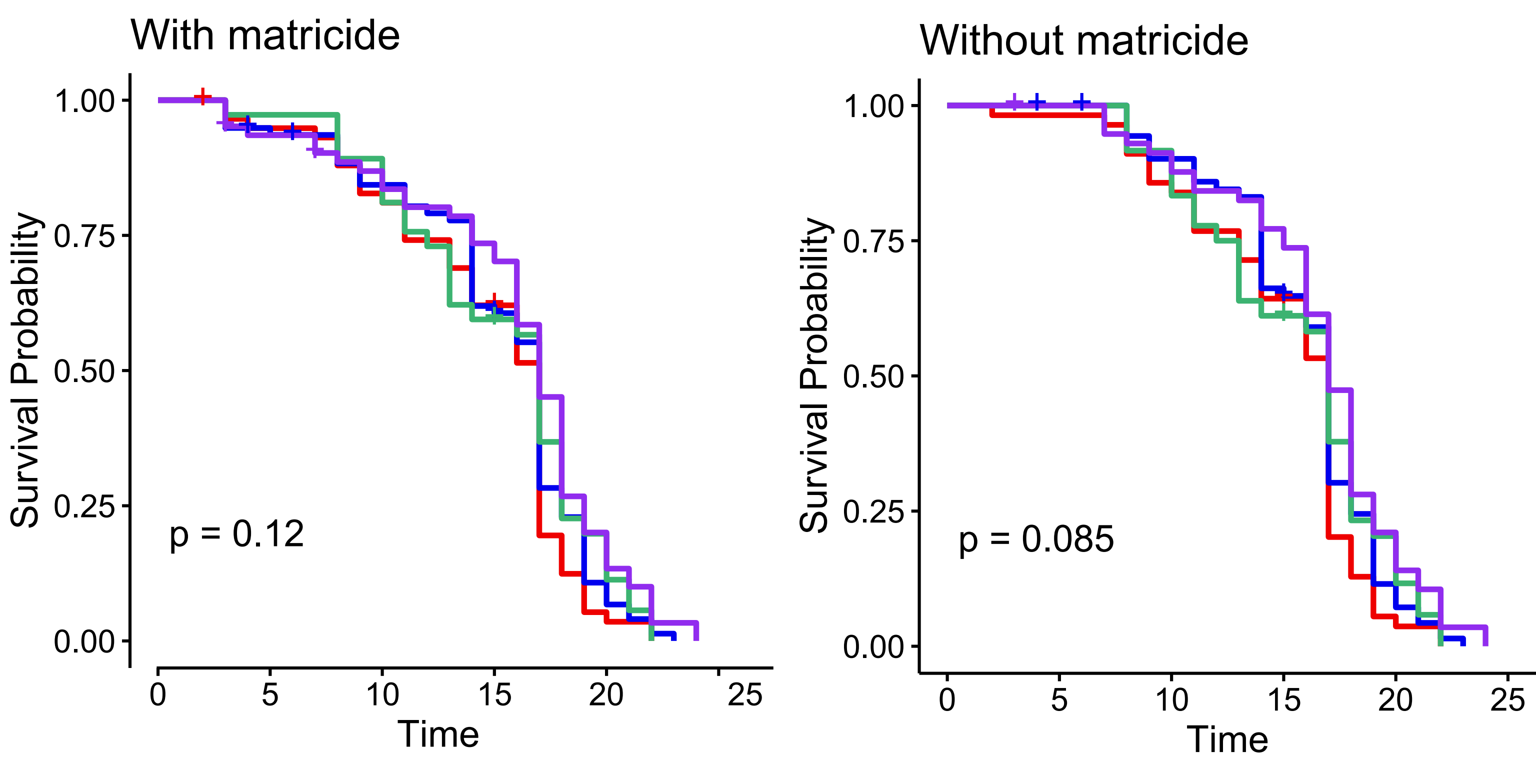

### Fig. S5

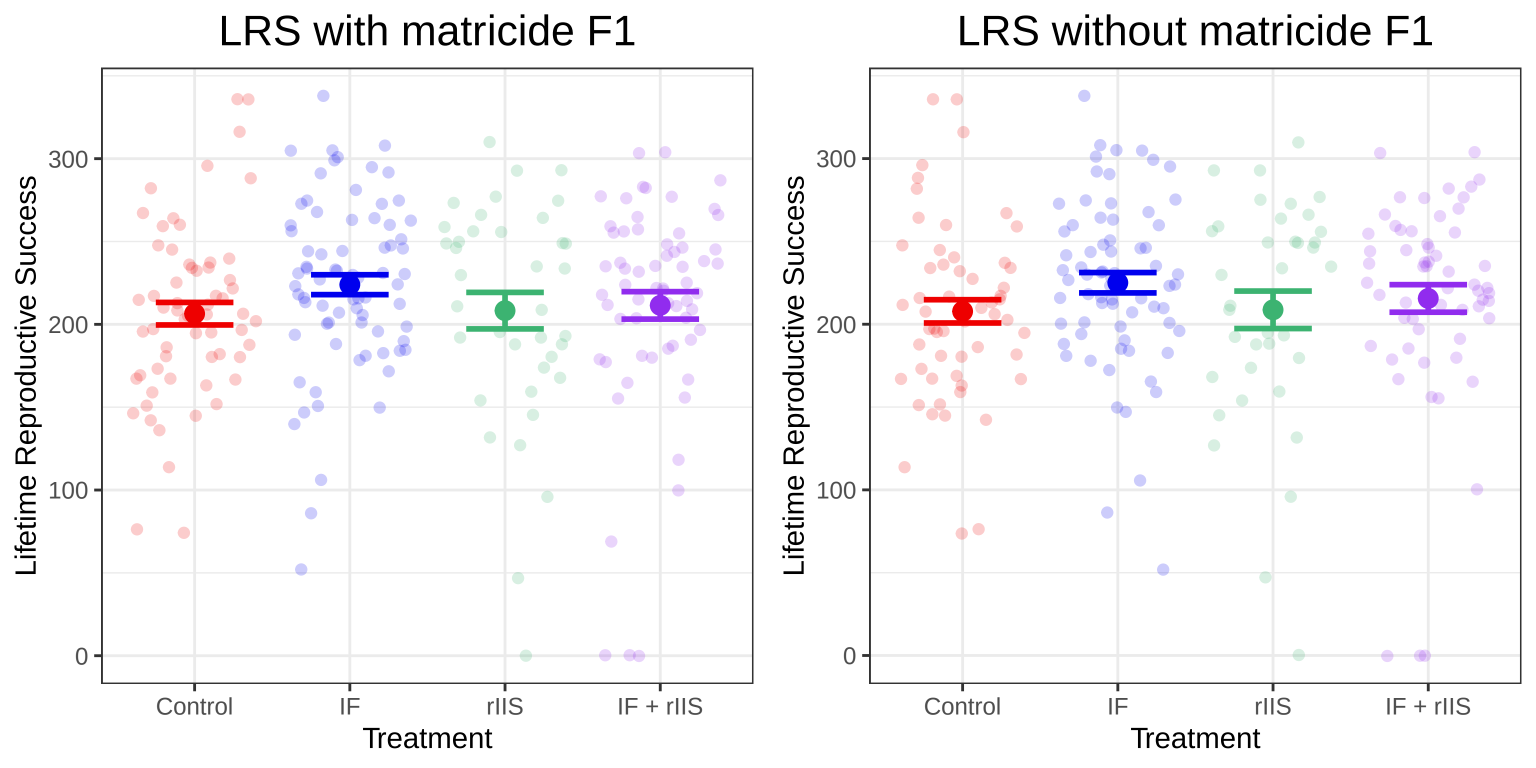

### Fig. S6

#### Principle Component Analysis

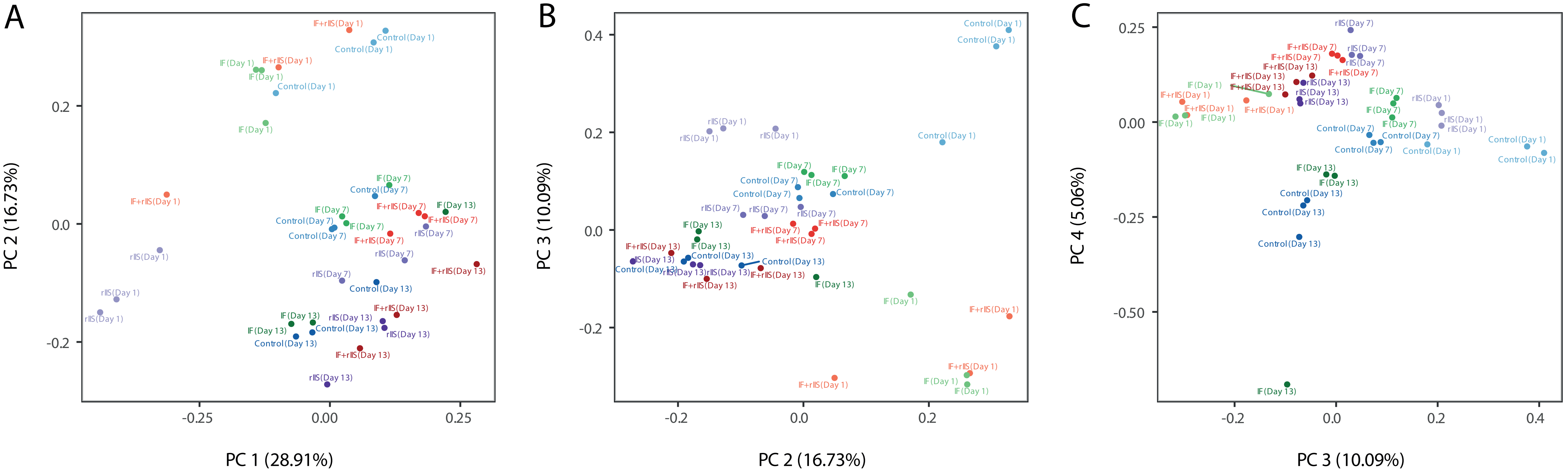

### Figs. S7 & 8

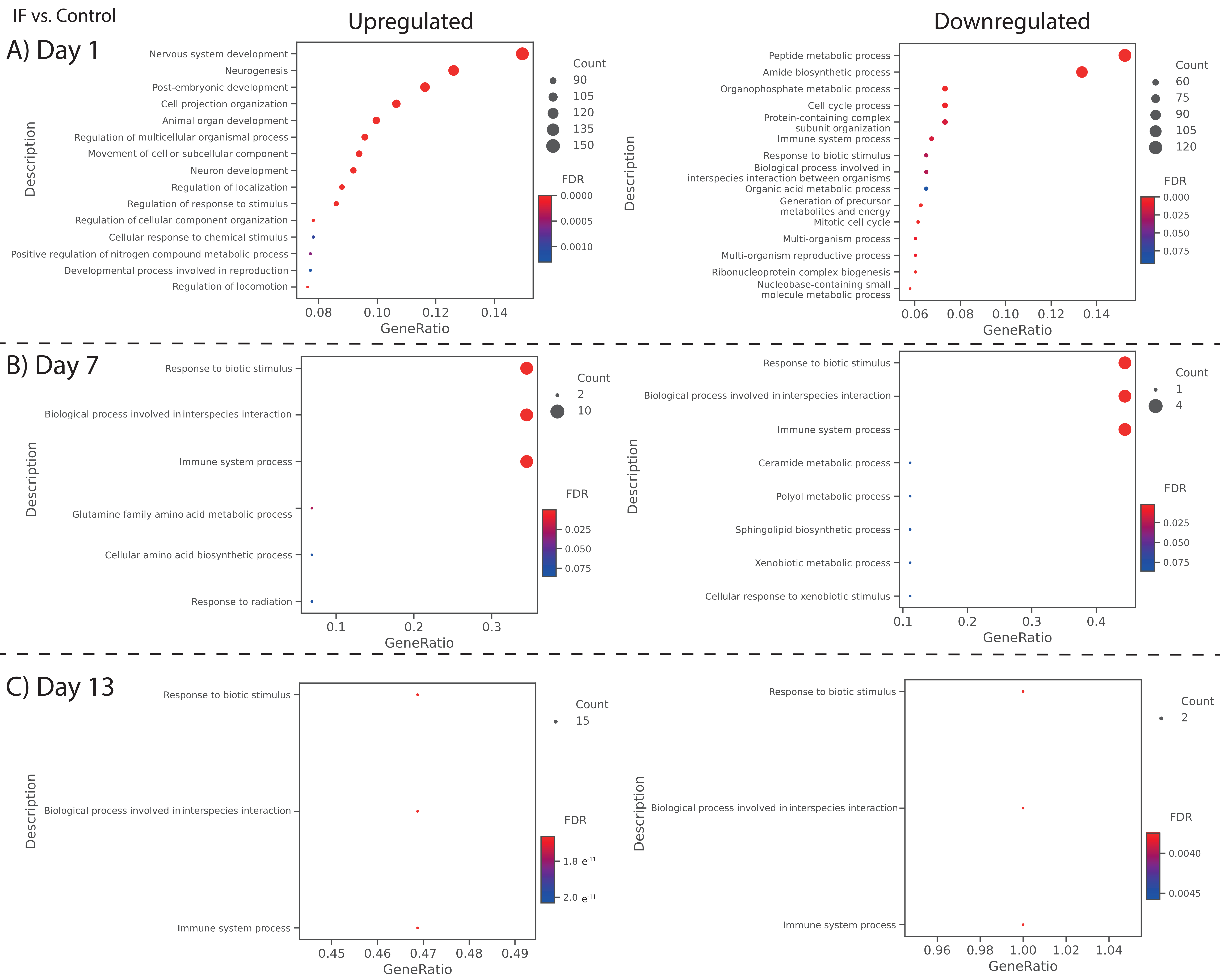

### Figs. S7 & 8

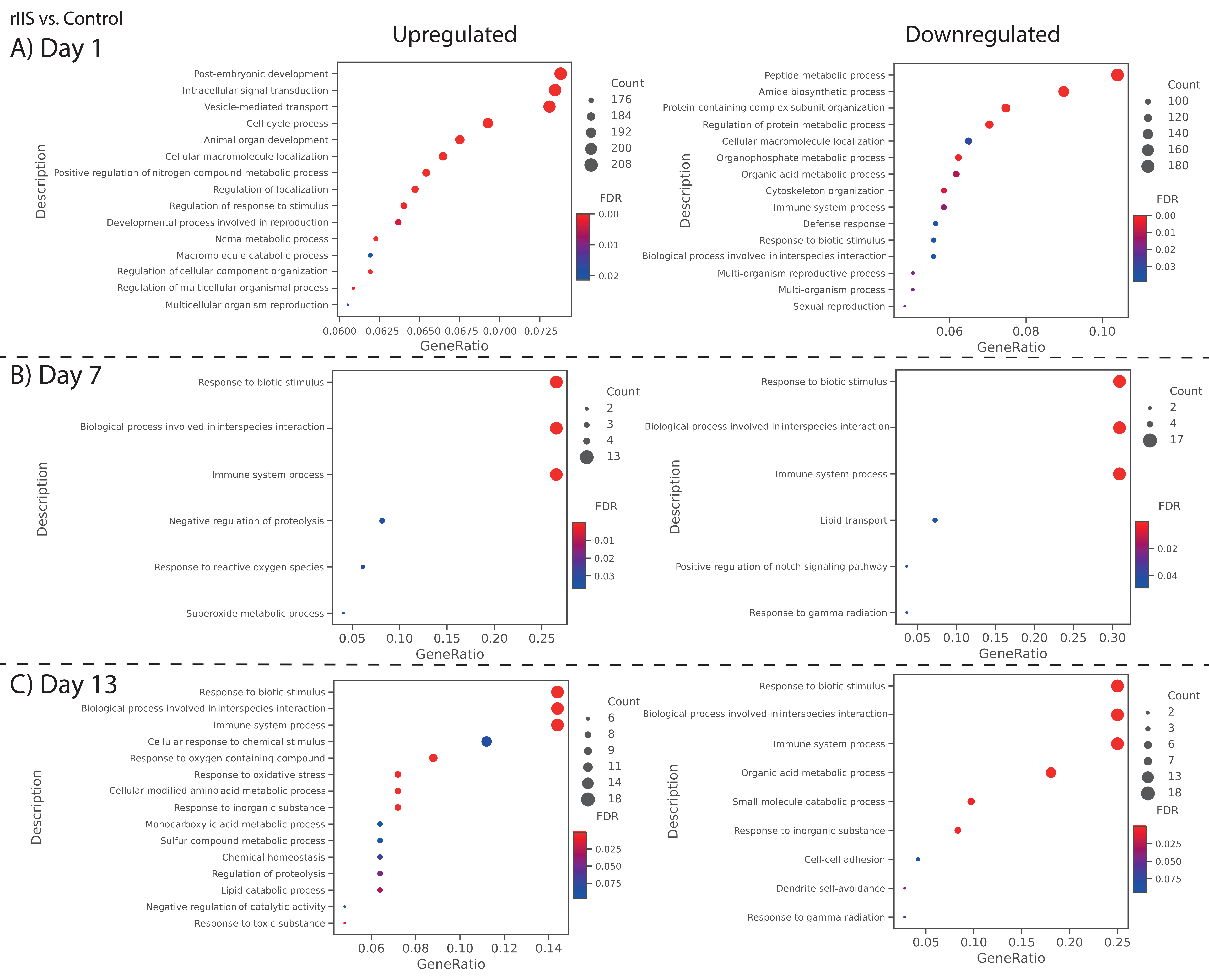
